# Alpha closed-loop auditory stimulation modulates waking alpha oscillations and sleep onset dynamics in a phase-dependent manner in humans

**DOI:** 10.1101/2022.11.15.512722

**Authors:** Henry Hebron, Beatrice Lugli, Radost Dimitrova, Valeria Jaramillo, Edward Rhodes, Nir Grossman, Derk-Jan Dijk, Ines R. Violante

## Abstract

Alpha oscillations play a vital role in managing the brain’s resources, inhibiting neural activity as a function of their phase and amplitude, and are changed in many brain disorders. Developing minimally invasive tools to modulate alpha activity and identifying the parameters that determine its response to exogenous modulators, is essential for the implementation of focussed interventions. We introduce Alpha Closed-Loop Auditory Stimulation (*αCLAS*) as an EEG-based method to augment and investigate these brain rhythms in humans with specificity and selectivity, using targeted auditory stimulation. Across three independent studies, we demonstrate that *αCLAS* alters alpha power, frequency, and connectivity in a phase, amplitude and topography-dependent manner. Using a single-pulse-*αCLAS* evoked potentials approach we show that the effects of auditory stimuli on alpha oscillations and resulting evoked potentials can be explained within the theoretical framework of oscillator theory and a phase-reset mechanism. Finally, we demonstrate the functional relevance of our approach by showing that *αCLAS* modulates sleep onset dynamics in an alpha phase-dependent manner.

## Introduction

The alpha rhythm (∼10 Hz) is a defining electrophysiological feature of the waking human brain, most prominent in parietal and occipital areas [1], but observed across a variety of neural regions [2,3]. Alpha oscillations have been associated with fundamental processes, from memory [4] and perception [5–8], to ageing [9,10] and disease [11–13]. Despite its reputed importance, selective modulation of the alpha rhythm remains challenging, and little is known about the underlying determinants of an alpha oscillation’s response to stimuli.

One promising route to efficacious alpha modulation is to address this rhythm on its own terms, using a closed-loop approach which leverages the instantaneous features of the oscillation in real-time. Influential theories posit that neuronal excitability fluctuates as a function of alpha phase and amplitude [14,15], since neurons more readily fire during the trough of alpha oscillations, and during periods of lower alpha amplitude [2,3,16,17]. Accordingly, the amenability of alpha to exogenous influence should itself also be phase-dependent, owing to the interdependent nature of spikes and oscillations [18].

In this vein, alpha phase-dependencies have been investigated in visual perception [5,8]. These studies typically sort the EEG, post-hoc, on the (binary) basis of perception of a visual stimulus, finding an effect of alpha phase at stimulus onset. However, phase estimates are often dependent on what proceeds them (i.e. they are non-causal) and conscious perception itself may have an electrophysiological component which backwards-influences the preceding phase estimate [19,20]. Experiments in which specific phases of the alpha oscillation are targeted in *real-time* are therefore needed to clarify phase-response dependencies.

Phase dependency of the response to stimuli has been demonstrated in a real-time manner for the transcranial magnetic stimulation (TMS) motor evoked potential (MEP) [21–24]. This approach circumvents the issue of time-dependency, but TMS pulses cause large artefacts in the EEG, rendering rapidly-repeating closed-loop (see [25]) stimulation unfeasible, and are clouded by concomitant residual auditory and somatosensory activity resulting from the TMS pulse [26].

Therefore, previous studies have not been able to provide neurophysiological characterisations of phase-dependencies of the alpha rhythm, i.e. whether alpha oscillations themselves respond to stimuli in a phase-dependent manner.

Closed-loop *sensory* stimulation has the potential to overcome this limitation, since sensory stimuli do not create artefacts in the EEG (allowing repeated neurophysiological interrogation), with the additional advantage of utilising safe, easy to apply and affordable technology. This approach has been used in sleep studies in which *sounds* are commonly phase-locked to the slow-oscillations (0.5-4Hz) of sleep [27–33]. Some of these studies have found a phase-dependent effect of stimuli and, in this case, their primary outcome is often neurophysiological. Thus, sounds boost or diminish slow-oscillations on a phase-dependent basis [27–33], although this remains contentious[29]. However, the closed-loop approaches used to deliver sounds at particular phases of the slow oscillation cannot be directly employed to target the much faster alpha oscillation, since they predominantly operate on the basis of an amplitude-threshold, which is unsuitable for alpha rhythms because they are highly variable in their amplitude and are instead defined on the basis of frequency. Recently, a number of strategies have been proposed to overcome this limitation, either relying on forecasting algorithms for real-time phase prediction [21] or through the application of a causal band-pass filter, the endpoint corrected Hilbert transform (ecHT) [34], to compute the instantaneous phase of an oscillatory signal in real-time.

Here, we introduce **Alpha Closed-Loop Auditory Stimulation (*αCLAS*)**, which uses the ecHT approach [34] to administer sound stimuli phased-locked to the alpha rhythms of the waking brain. Across three independent studies we employ *αCLAS* to explore the precise contributions of *phase* to the response of alpha rhythms to external stimuli and provide evidence of the functional implications of this approach. We first demonstrate that repetitive-*αCLAS* induces phase-dependent modulations of alpha power, frequency, and connectivity in a spatially-specific manner. We then use single-pulse *αCLAS* to show that these effects are dependent on oscillation amplitude and can be explained using oscillator theory and a phase-reset model. Finally, we apply these principles during the transition to sleep, a process in which alpha oscillations become naturally dampened, to show that the dynamics of sleep onset can be modulated in a phase-dependent way by *αCLAS*.

## RESULTS

### Study 1. Repetitive-αCLAS – Proof of Principle

#### Accuracy and spatial specificity of alpha closed-loop auditory stimulation

In Study 1, we examined whether repetitive-*αCLAS* can selectively modulate the resting alpha oscillation at different cortical locations, on the basis of the phase at which sounds are repeatedly administered (at four orthogonal phases: pre-peak 330°, post-peak 60°, pre-trough 150°, post-peak 240°, **Figure 1A**). Phase was computed in real-time (using the endpoint-corrected Hilbert Transform, ecHT [34]) based on the EEG activity recorded from one electrode placed over the frontal, Fz (experiment 1) or parietal cortex, Pz (experiment 2). Simultaneously, we recorded whole-scalp activity using high-density (hd) EEG. Participants were exposed to 30-seconds of repeated sounds (termed ‘on’), interleaved with 10-seconds of silence (‘off’), while seated with their eyes closed (experiment 1, N=20, experiment 2, N=28; 10 blocks per phase per experiment, **Figure 1B**).

**Figure 1.**
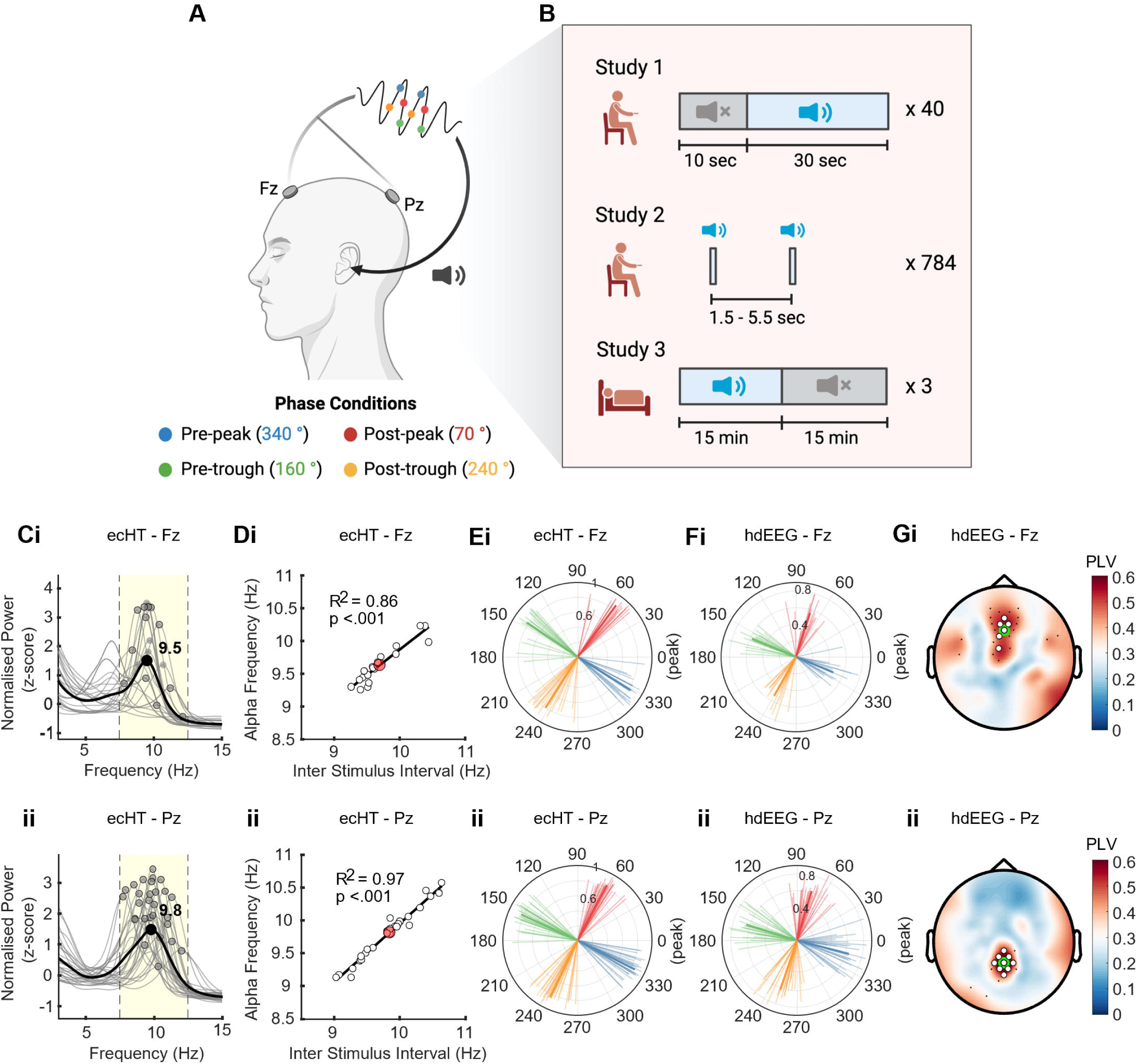
Study Overview and Phase-locking validation. (**A**) Schematic of closed-loop auditory stimulation. The signal is processed in real-time to compute its instantaneous phase (coloured dots correspond to the 4 orthogonal phases targeted), and pulses of pink noise (20 ms duration, 80 dB volume) are delivered at the pre-specified phase of the target rhythm. Phase information is extracted from the EEG signal at frontal, Fz, or parietal, Pz, cortical locations (Fz and Pz refer to positions on the 10-20 EEG International system). **(B)** Pictorial summary of the 3 studies conducted, indicating periods of sound ‘on’ stimulation and ‘off’ periods, as well as the number of trials per phase condition. **(C)** Normalised (z-score) power spectrum computed for ‘off’ periods, taken from the phase-locking (i.e. ecHT) electrode in **(i)** experiment 1 (Fz) and **(ii)** experiment 2 (Pz). Phase-locked band (7.5-12.5 Hz) highlighted in yellow. **(D)** Stimulation (Inter-Stimulus Interval; ISI) vs Alpha frequency as computed for ‘off’ periods only, taken from the phase-locking electrode in **(i)** experiment 1 (Fz) and **(ii)** experiment 2 (Pz). White circles represent individual participants, red circle shows group average. **(E)** Average phase angle and resultant at stimulus onset taken from the phase-locking electrode in **(i)** experiment 1 (Fz) and **(ii)** experiment 2 (Pz). Thin lines represent individual participants, thick lines show the group average. **(F)** Same as **(E)** but computed using the respective electrodes from the high-density (hd) EEG; **(i)** experiment 1 (Fz) and **(ii)** experiment 2 (Pz). **(G)** Topoplot of resultant - shows the mean resultant vector length for each phase condition at all channels of the hd-EEG data. White marks indicate channels at which resultant >0.5 and p <0.05. Black marks indicate channels at which resultant <0.5 and p <0.05. Green circles indicate the position of the target electrode. p values from Bonferroni-corrected z-test for non-uniformity. Shown for **(i)** experiment 1 and **(ii)** experiment 2.

We first examined whether sounds were phase-locked in a physiologically meaningful manner. The presence of a spectral peak in the alpha band (7.5-12.5 Hz) in the phase-locking electrode during the ‘off’ period was confirmed in both the group average and majority of participants (experiment 1: 18/20; experiment 2: 28/28 participants, **Figure 1C**). The individual alpha frequency (IAF) in the ‘off’ periods correlated strongly with the average inter-stimulus interval (ISI) of the sounds, indicating that stimulation frequency was tailored to the individual, with approximately 1 stimulus applied per alpha cycle (**Figure 1D**).

Phase-locking accuracy was then assessed by calculating the mean resultant vector length and average angle of phase at stimulus onset, resulting in a value between 0 (uniform distribution) and 1 (unimodal distribution, perfect phase-locking). The resultants were high for both experiments and for all phases, with significant unimodal distributions (experiment 1, resultant mean ± std: 0.84 ± 0.07; experiment 2: 0.85 ± 0.05; Z-tests confirm significant unimodality in all conditions; z values >18, ps<0.001, **Figure 1E**). Furthermore, the average phase angles were highly consistent across participants, and orthogonal between conditions (experiment 1, phase angle deviation from target, mean ± std: 3.80° ± 2.14; experiment 2: 3.63° ± 1.37). Likewise, resultants were high in the hd-EEG system (experiment 1, mean ± std: 0.54 ± 0.13; experiment 2: 0.63 ± 0.13; z values >8, ps<0.001, **Figure 1F**) and showed high phase-specificity. These results show that sounds were accurately delivered at the prescribed phase. Stimulus onset phase angles were comparable across experiments and EEG systems, but resultant vector lengths were lower in the hd-EEG, which can be explained by the difference in reference-schemes; the ecHT system used a mastoid reference, whilst the hd-EEG used scalp current density. Finally, we took advantage of the spatial resolution of the hd-EEG to assess the spatial selectivity of the phase-locking. Phase-locking was highly accurate across experiments, greatest around locations targeted (**Figure 1F** and **Figure S1**).

#### αCLAS induces phase and location-specific effects on alpha power and frequency

Having established the accuracy and spatial-specificity of *αCLAS*, we examined whether stimulation induced phase-dependent changes to the alpha oscillations we set out to target. When sounds were phase-locked to Fz EEG (experiment 1) we observed a main effect of phase on power, which was localised to frontal midline regions, around the target electrode (**Figure 2A** shows the topography of the effect of *αCLAS* phase determined by the ANOVA, for the centre, i.e. 10 Hz, alpha frequency; **Figures S2, S3** show the specificity of this effect in the alpha band).

**Figure 2.**
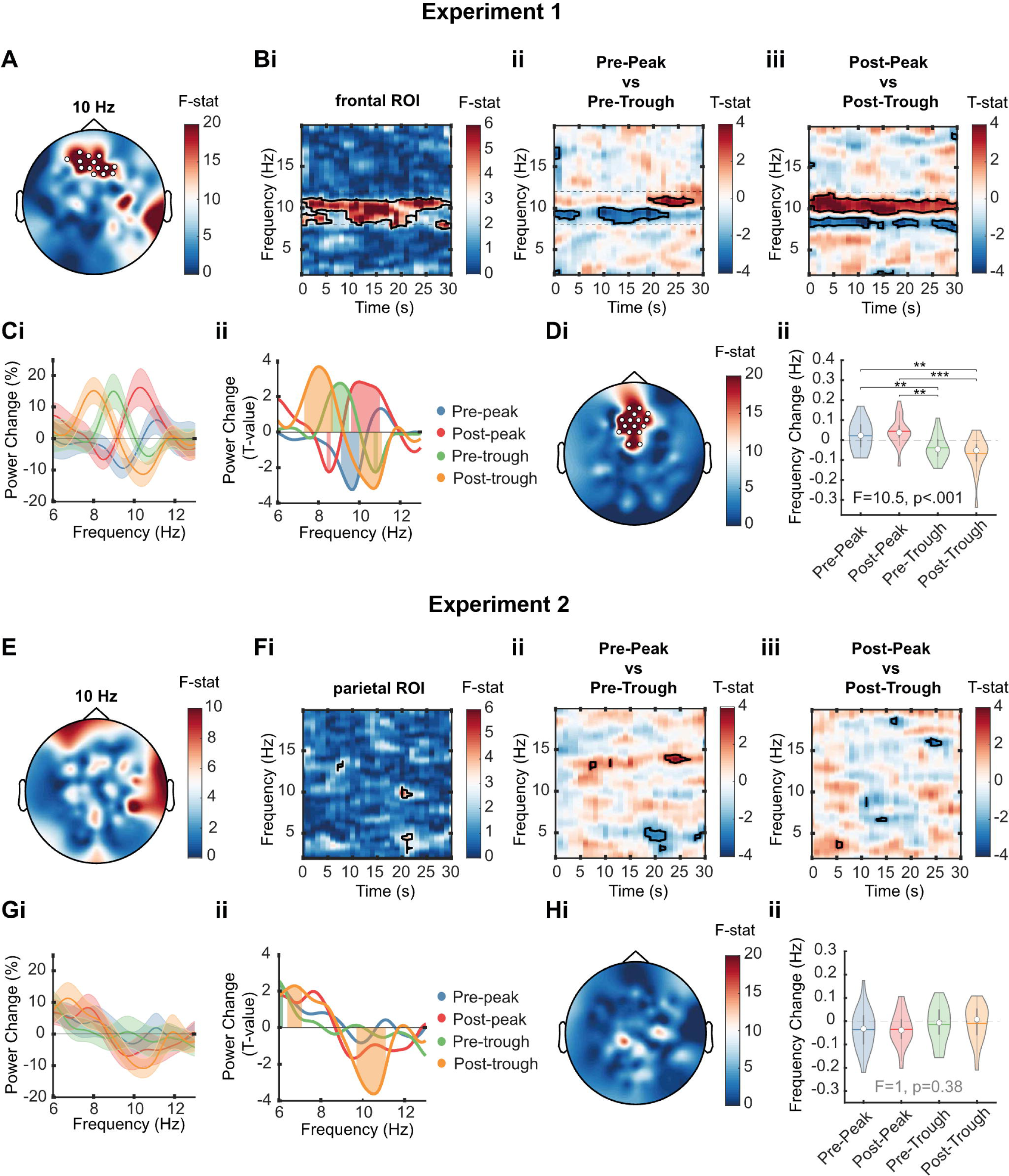
Stimulation-induced power and frequency changes (Study 1) **(Ai)** Topography of 10 Hz power change ANOVA. We computed the power spectral density for every phase condition in the ‘on’ and ‘off’ periods and expressed stimulation-induced changes in power as percentage change from ‘off’ to ‘on’ per frequency bin. White marks indicate cluster-corrected p<0.05. **(Aii)** Time-frequency ANOVA. Outlined clusters p < 0.05. **(Bi)** Time-frequency, pre-peak vs pre-trough, for the frontal region-of-interest (see ROI specification in Materials and Methods). Positive values indicate pre-peak > pre-trough. Outlined clusters ANOVA p < 0.05. **(Bii)** Time-frequency, pre-peak vs pre-trough, at respective regions of interest. Positive values indicate post-peak > post-trough. Outlined clusters t-test p < 0.05. **(Ci)** Power change spectrum (% change from ‘off’ periods) across the alpha-band frequencies for the four targeted phases collapsed across the stimulation period in the frontal ROI. **(Cii)** Power change spectrum (T-values, one-sample T-test) at respective region of interest. Shaded regions p<0.05. **(Di)** Topography of frequency change ANOVA. White marks indicate cluster-corrected p<0.05. **(Dii)** Frequency change violin plots, stats indicate linear mixed effects model results and contrasts, *** *p* < 0.001, ** *p* < 0.01, * *p* < 0.05, † *p* < 0.1, t-tests. **E-H** show the same analyses/results as **A-D**, but for experiment 2.

Changes in power were circumscribed to the alpha band and persisted throughout the 30-second stimulation period (**Figure 2Bi**). To better understand this effect, we compared power changes between opposite phases (**Figure 2Bii-iii**). We again observed significant effects of *αCLAS* phase in the alpha band that were present throughout the stimulation period, particularly when comparing the post-peak and post-trough conditions (**Figure 2Biii**). Noteworthily, these effects appeared to be frequency-specific, i.e., frequencies within the alpha band were not modulated equally. When different phases were targeted, the power at different frequencies within the alpha band was indeed significantly increased or decreased (**Figure 2C**). The manner in which these power changes were distributed suggested a slowing or quickening of alpha rhythms; in the post-trough condition, for example, power decreased for high alpha frequencies, and increased for low alpha frequencies (i.e. slowing), and the inverse was true for the opposite phase, i.e. post-peak. For pre-peak and pre-trough conditions frequency effects were more modest, with the former mostly decreasing power in the alpha band and the latter largely increasing it.

To verify whether different phases impacted the frequency of alpha oscillations differently, we estimated instantaneous frequency (within the alpha band) across all EEG channels, following the method proposed by Cohen (2014). The stimulation-induced frequency change for each condition was then computed by subtracting the average alpha frequency of ‘off’ from ‘on’ periods. The topography of the effect of phase on frequency (**Figure 2Di**) showed a strikingly similar distribution to the effect of phase on power (**Figure 2A**). Comparison of the frequency change showed significant differences in frequency between each phase condition, in agreement with what was inferred from the power change spectra, i.e. a slowing or quickening of different extents (**Figure 2Dii**, these differences were not present during the ‘off’ period, **Figure S4**).

We then performed the same analyses to experiment 2, targeting Pz. Contrary to our previous results, we did not observe a significant effect of *αCLAS* phase on the power of alpha oscillations (**Figure 2E-F** and **Figures S5**, **S6**). We observed a significant modulation of alpha power when focusing on a parietal ROI centred around Pz, but only for the post-trough, where alpha frequencies between ∼10-11 Hz were reduced (**Figure 2G**). However, we did not detect a change in frequency for Pz stimulation (**Figure 2H** and **Figure S7**). Overall, these results demonstrate that the effects of *αCLAS* on alpha power and frequency are phase and location specific.

#### αCLAS induces whole-brain changes in connectivity depending on the phase and targeted location

Functional connectivity metrics provide a useful framework to understand the coordination of brain regions during specific states [36,37]. We evaluated whether sounds altered brain connectivity in a phase-dependent manner by using two well-established and complementary, phase-based measures of synchrony, the phase-locking value (PLV) and phase lag index (PLI). While PLV is more sensitive to type 1 errors, i.e. incorrectly assigning connectivity (rejecting the null) that is in fact volume conduction, PLI is more sensitive to type 2 errors, i.e. incorrectly rejecting genuine connectivity (failing to reject the null) with zero (modulo pi) phase difference.

We observed a significant effect of *αCLAS* phase on the connectivity around the phase-locking site when targeting the alpha rhythm at Fz, experiment 1 (**Figure 3Ai**), which was specific to the alpha band (**Figure S8**). To explore whether this frontal midline area modulated its connectivity to other regions of the brain, we used the electrodes showing significant phase effects as seeds and computed their connectivity to all EEG channels. The connectivity of this frontal cluster differed primarily between phase conditions in short-range connections with itself, and long-range connections with occipito-parietal and central channels (**Figure 3Aii**). To better understand the effect of *αCLAS* phase on connectivity we computed PLV per phase condition (**Figure 3B**). The most noticeable patterns were observed for the post-peak and post-through conditions, such that during post-peak stimulation there was an increase in connectivity, particularly between frontal midline and occipito-parietal channels, in good agreement with the topology observed in **Figure 3Aii**. The post-through condition, on the other hand, resulted in a fairly global decrease in connectivity, which was also present, albeit to a smaller extent, for the pre-peak and pre-through conditions. Using PLI, we observed similar effects of phase on connectivity, which was centred around Fz (**Figure 3Ci**), and comparable patterns to PLV of local and long-range connectivity between the frontal midline cluster and occipito-parietal channels (**Figure 3Cii**). Regarding PLI per phase condition, while the overall pattern was similar to PLV, long-range connectivity was less salient, suggesting either the presence of volume conduction or the true existence of zero modulo pi-lag connectivity between these regions (**Figure 3D**).

**Figure 3.**
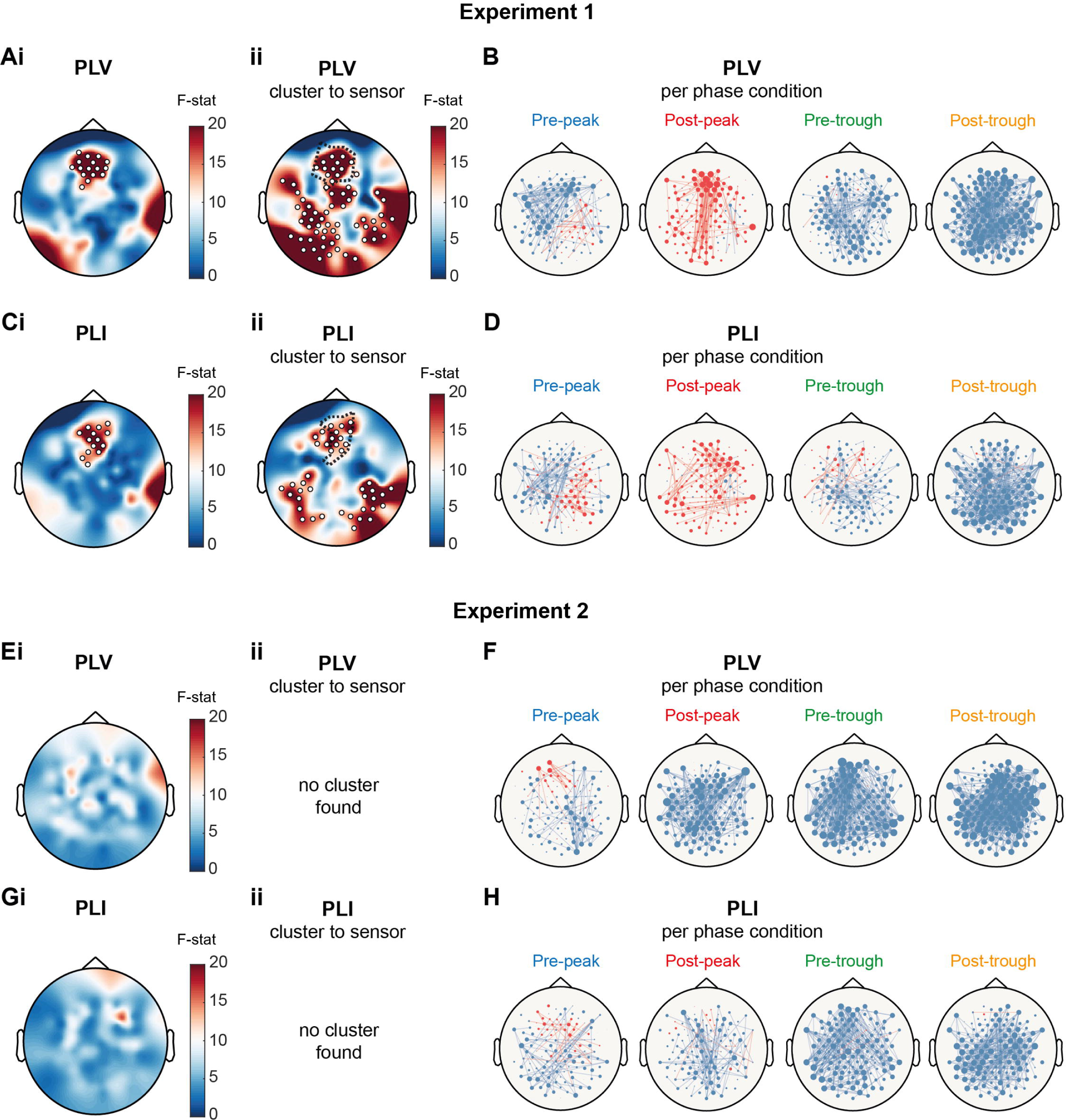
Stimulation-induced connectivity changes in alpha band (Study 1) **(Ai)/(Ei)** Topography of main effect of phase on average alpha band PLV for each channel as per ANOVA. **(Aii)/(Eii)** Topography of average alpha band PLV for channels identified in cluster (dotted line shows identified cluster).* **(Ci)/(Gi)** Topography of variance between conditions of average alpha band PLI for each channel as per ANOVA.* **(Cii)/(Gii)** Topography of average alpha band PLI for channels identified in cluster (dotted line shows identified cluster).* **(B)/(F)** Stimulation-induced changes in alpha band PLV per condition as per t-test (compared to the ‘off’ period), lines are plotted where *p*<0.01. Red lines indicate an increase in the connectivity of that channel pair, blue lines indicate a decrease. **(D)/(H)** Stimulation-induced changes in alpha band PLI per condition as per t-test, lines are plotted where *p*<0.01. Red lines indicate an increase in the connectivity of that channel pair, blue lines indicate a decrease. *White marks indicate cluster-corrected p<0.05

Analysis of connectivity for experiment 2 (i.e. targeting Pz), did not show an effect of phase on either alpha PLV (**Figure 3E**) or alpha PLI (**Figure 3G**), although a small and disparate effect was seen for delta PLV and beta PLI (**Figure S9**). Exploratory analysis on the alpha connectivity patterns of each phase condition showed a generalised decrease in whole-brain connectivity, particularly for pre-through and post-through conditions, in both PLV (**Figure 3F**), and PLI (**Figure 3H**).

Overall, we observed again that the effects of *αCLAS* on brain dynamics are dependent on the phase applied and the location of the alpha rhythm targeted.

#### Changes in connectivity are related to frequency changes induced by αCLAS

Changes in connectivity might be explained by differences in frequency; in that the synchrony between two oscillators will, at least partially, depend on the differential of their frequencies – it is not possible for oscillators of different frequencies to have a coupling constant of 1, for example. Accordingly, we explored whether connectivity and frequency were related in experiment 1, where phase-dependent changes in frequency and connectivity were observed.

We first investigated whether frequency differences are present across the scalp in the absence of stimulation. This was achieved by expressing alpha frequency at all channels relative to frontal ROI alpha frequency, before averaging these values across all ‘off’ blocks of all conditions and participants to produce **Figure 4A**. On average, alpha frequency was lowest at the frontal ROI and highest (approximately 0.4 Hz higher) across parietal and occipital regions. This gradient is consistent with previous investigations of frequency [38]. **Figure 4B** shows that when comparing the absolute frequency at the frontal ROI to all remaining channels, the frontal ROI was slower across all but one participant. This means that a selective increase in alpha frequency at the frontal ROI would bring it more in line with the remainder of the scalp, and a decrease would do the opposite. We then calculated the stimulation-induced change in this frequency difference by subtracting the absolute frequency difference in ‘on’ periods from those of ‘off’ periods for each condition; meaning regions becoming more similar in their frequencies with stimulation are assigned a negative value and vice versa. **Figure 4C** shows that the frequency difference was changed in an *αCLAS* phase-dependent manner, in accordance with the previously outlined frequency changes in **Figure 2**.

**Figure 4.**
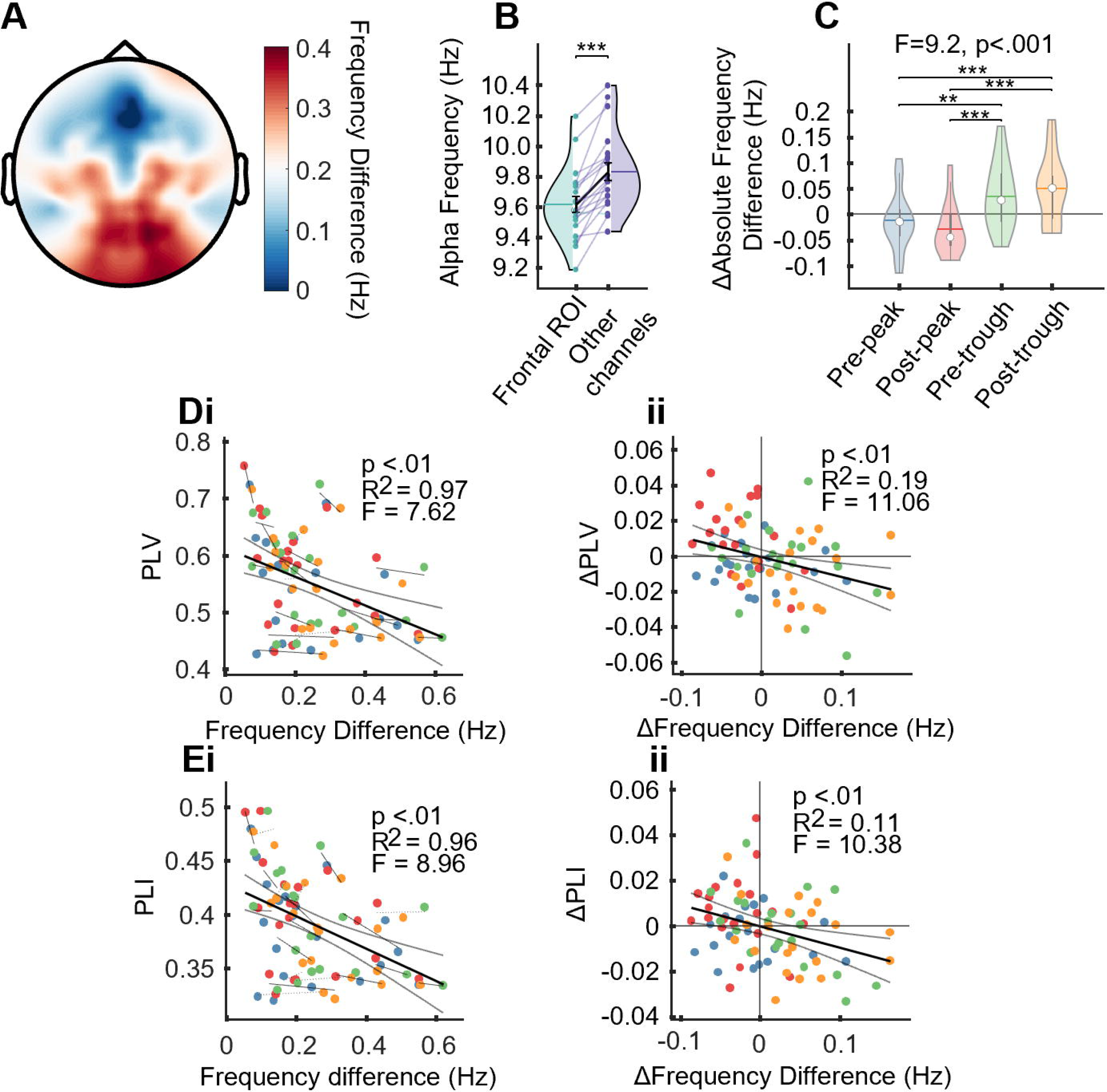
Connectivity is partially accounted for by frequency difference in experiment 1. **(A)** Topography of alpha frequency difference from frontal ROI during ‘off’ (i.e. no stimulation) periods. **(B)** Alpha frequency at frontal ROI vs all other channels during off periods.^†^ **(C)** Average stimulation-induced changes in alpha frequency differences, between frontal ROI and all other channels. ^††^ **(D) (i)** Average frequency difference between frontal ROI and all other channels vs average PLV between frontal ROI and all other channels, lines indicate individual participant linear fit across four conditions^†††^ **(ii)** Average stimulation-induced frequency difference change between frontal ROI and all other channels vs average stimulation-induced PLV change between frontal ROI and all other channels. Crossed lines indicate zero.^†††^ **(E)(i)(ii)** same as **(D)** but for PLI^†††^. ^†^Stats are from a paired t-test, *** *p*<0.001 ^††^Stats are from a LMEM: ΔAbsolute_frequency_difference ∼ Condition + (1|Participant) ^†††^ Stats are from a LMEM; Connectivity ∼ Frequency_difference + (1|Participant) or ΔConnectivity ∼ ΔFrequency_difference + (1|Participant)

Finally, we quantified the relationship between frequency difference and connectivity using all data from all conditions in a mixed effects model with participant as a random effect. This showed the predicted significant negative relationship between absolute frequency difference and connectivity in both of our connectivity metrics, and this held true for both the absolute values (**Figure 4Di** for PLV and **Figure 4Ei** for PLI) and the stimulation-induced changes to those values (**Figure 4Dii 4Eii**). Although changes in frequency difference explain only a small amount of variance in connectivity (PLV R^2^=0.19, PLI R^2^=0.11), we suggest that it is a contributory factor and may, at the very least, describe differences in connectivity between the frontal ROI and the remainder of the scalp. This also indicates that a genuine change in connectivity may be at play, in line with theories of intra-brain information transfer [14,39,40]; if inter-areal communication is dependent on oscillation phase, there may be a particular phase at which sound-evoked activity is most prominently relayed between regions.

### Study 2. Single-Pulse αCLAS – Mechanism

#### Auditory evoked potentials provide a putative phase-reset mechanism by which αCLAS may operate

To understand why sounds administered at different phases of the alpha oscillation appear to speed it up or slow it down we conducted a new study similar to Study 1, but with the critical difference that sounds were administered as isolated/single-pulse stimuli. In addition to allowing an exploration of phase reset, the application of single-pulse *αCLAS* provides a means to further investigate the spatial specificity (Fz vs Pz stimulation) of the effects reported before. In a phase-reset model, the phase (and hence, the *frequency*) of an oscillation changes *abruptly* following reception of a stimulus. Specifically, this model predicts that the extent and trajectory (i.e. faster or slower) of the reset is dependent on the phase at the onset of perturbation, consistent with our findings from experiment 1. For example, if an oscillation reliably resets to a particular phase with a fixed latency following a stimulus, the trajectory of its phase (and consequently, the frequency of the oscillation) would differ depending on the phase at stimulus onset – this can take two forms: an advancing of the cycle or a delaying, to meet the reset [41].

Study 2 was composed of two experiments targeting the alpha rhythm at Fz and Pz, respectively (experiment 3: N=8; experiment 4: N=7). Participants sat with their eyes closed and listened to single pulses of sound (196 per phase condition), locked to the same 4 phases as before, and interleaved with silent periods, **Figure 1B**.

We first assessed the profile of the auditory-evoked potential (AEP) responses following sound stimulation to each phase condition. We observed the stereotypical AEP response, more pronounced in central and frontal than parietal brain areas, e.g.[42,43], (**Figure 5Ai**, **Figure 5Fi**). Before sound onset (at time 0) the four *αCLAS* phase conditions showed, as expected, orthogonal timecourse profiles (ANOVA shows a main effect of phase on signal amplitude). The pre-stimulus amplitude was also higher at Pz (**Figure 5Fi**) in agreement with the more pronounced distribution of alpha in occipito-parietal areas at rest with eyes closed [1] (mean pre-stimulus log alpha band power ± std, experiment 3: -6.30 ± 0.38; experiment 4: -5.58 ± 0.60; two sample t-test: *t*(14) = -2.67, p = 0.018). The post-stimulus response shows a high overlap between conditions at Fz, whilst at Pz there remained significant variance between conditions even following stimulus onset.

**Figure 5.**
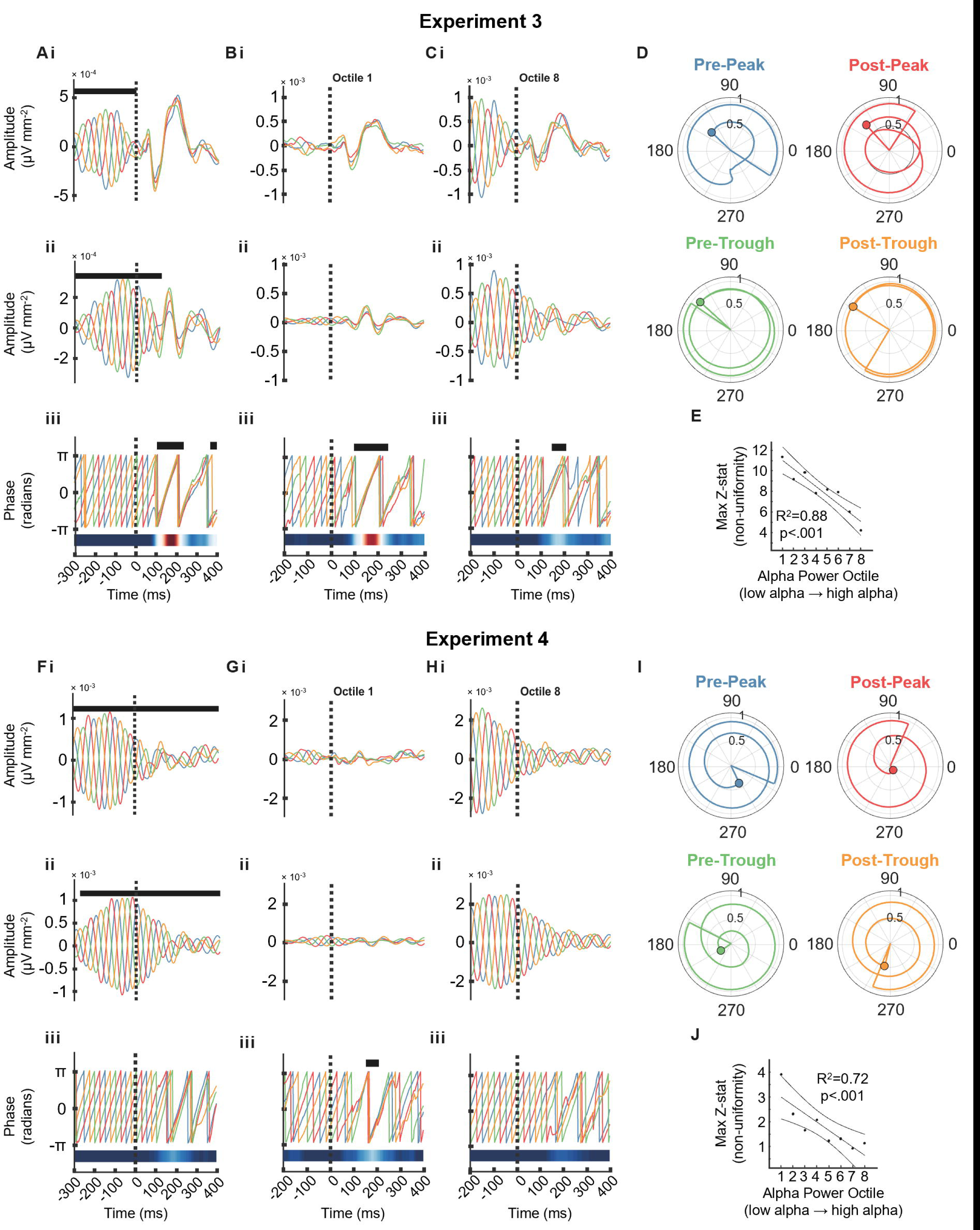
Auditory evoked potential (AEP) and Phase Reset (Study 2) **(A and F)** AEP at Fz in experiment 3 (**A**) and Pz in experiment 4 (**F**). (**B** and **G**) Lowest pre-stimulus alpha power octile at Fz in experiment 3 (**B**) and Pz in experiment 4 (**G**). (**C** and **H**) Highest pre-stimulus alpha power octile at Fz in experiment 3 (**C**) and Pz in experiment 4 (**H**). (**A, B, C, F, G, H**) **i.** Broadband (1-40 Hz) AEP*. **ii.** Amplitude component of alpha band (7.5-12.5 Hz) endpoint-corrected Hilbert transformed AEP*. **iii.** Instantaneous phase of alpha band (7.5-12.5 Hz) from endpoint-corrected Hilbert transformed AEP**. **(D** and **I**) Phase trajectory in circular space for each of four phase conditions across first 200 ms at Fz in experiment 3 **(D)** and Pz in experiment 4 (**I**). Vector without circle indicates average phase and resultant of the group at time 0, vector with circle indicates average phase and resultant of the group at time 200 ms. Circular line shows trajectory (in phase and resultant of the group) between the two timepoints. **(E** and **J**) Pre-stimulus alpha power vs maximum post-stimulus Z-statistic (i.e. phase synchrony across conditions) at Fz in experiment 3 (**E**) and Pz in experiment 4 (**J**), line, confidence intervals and statistics from linear regression. * Black marks indicate ANOVA *p* <0.05 ** Black marks indicate Rayleigh test *p* <0.05, heatmap shows time series of Z-statistic

The trajectory of both the amplitude (**Figure 5Aii**, **Figure 5Fii**) and phase (**Figure 5Aiii**, **Figure 5Fiii**) components of alpha, explored using the ecHT algorithm offline, showed once more the orthogonality of conditions prior to stimulation and their striking uniformity following stimulation (Rayleigh test at each time point, confirming that the conditions became significantly aligned at Fz in experiment 3 but not in experiment 4). Circular plots, showed that in experiment 3 the phases became aligned between conditions over 200 ms, and the resultants remained high (**Figure 5D**), whilst this was not the case for experiment 4 (**Figure 5I**).

To address the effect of oscillation strength on perturbability, we sorted the AEPs into eight octiles based on pre-stimulus alpha amplitude. At Fz, the magnitude of the broadband AEP was comparable between the lowest pre-stimulus alpha power octile (**Figure 5Bi-ii**) and the highest one (**Figure 5Ci-ii**), despite clear differences in the pre-stimulus activity, suggesting that the two components are not strongly dependent.

However, when looking at the alpha phase component (**Figure 5Biii**, **Figure 5Ciii**) of the AEP, the lowest pre-stimulus alpha power octile showed stronger synchrony, with longer periods of unimodality. Indeed, even at Pz there was post-stimuli phase alignment when the pre-stimulus alpha power was at its lowest amplitude (**Figure 5Giii**). Moreover, we observed a strong linear relationship between alpha octile and reset magnitude at both Fz and Pz (**Figure 5E**, **Figure 5J**).

#### Further Characterisation of the Phase Response to Alpha Phase-Locked Sound Perturbations

A phase-reset is often explored using phase transfer curves (PTC’s) [41,44–46], in which the starting phase (i.e. at stimulus onset) is plotted against end phase (i.e. phase after a certain amount of time has expired). Phase response curves (PRC’s) provide similar information, but instead of the end phase, the phase change is plotted. These curves allow both the magnitude and the specific nature of the reset to be summarised, detailing which start phases lead to an advance or delay of the cycle, and to what extent.

The PRC’s and PTC’s confirmed that sounds phase-locked at Fz, in experiment 3 (**Figure 6A-C**), induced a greater reset by 200 ms than Pz in experiment 4 (**Figure 6D**-**E**), although there may still be a weak reset at Pz as previously observed (**Figure 5Giii**). Oscillator theory would distinguish these two types of resetting as type 0 and type 1 respectively – type 0 describes a strong input/reset, characterised by a phase response curve with a gradient of 0 (as in Fz, **Figure 6A**), whilst type 1 describes a weak input/reset, typified by a phase response curve with a gradient of 1 (as in Pz, **Figure 6D**) [41,44–46]. Additionally, when targeting Fz, the phase-dependent slowing and quickening observed in the PRC, agrees with what was seen in experiment 1; post-peak resulted in a quickening and post-trough achieved the opposite, but the remaining two conditions show more ambiguous responses. This ambiguity may be explained by the complexity associated with repeated perturbations. If an oscillator phase-resets with a fixed latency following a perturbation, but its frequency is allowed to vary, the phase-response profile should be frequency dependent – thus, if one pulse brings about a change in frequency, a second pulse may interact with the oscillation differently.

**Figure 6.**
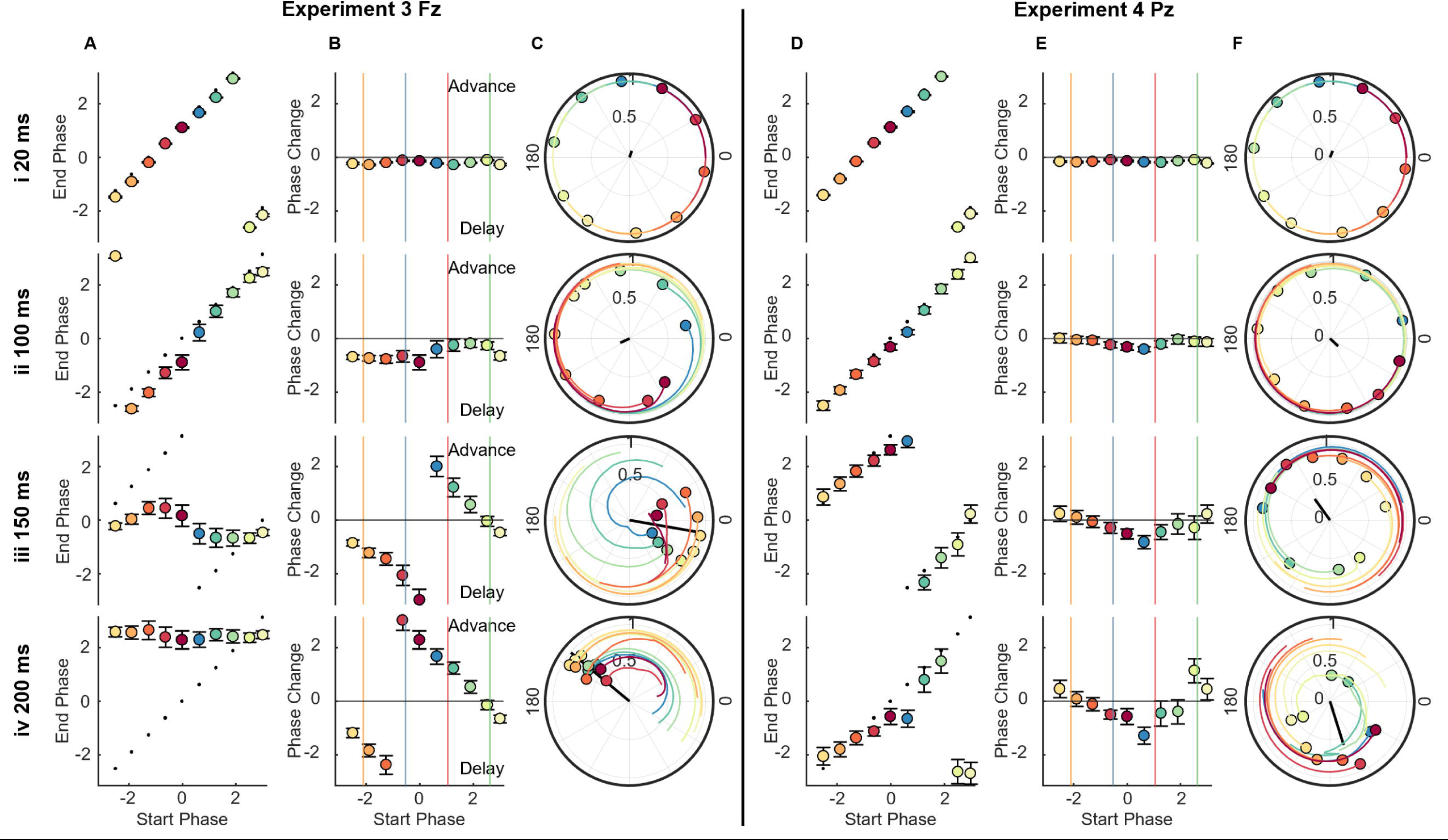
Phase Transfer and Phase Response Curves (Study 2) **(A and D)** Phase Transfer Curve, showing start phase vs end phase (in radians) for Fz in experiment 3 (**A**) and Pz in experiment 4 (**D**). Dashed line indicates predicted phases, based on a 10 Hz oscillation. Error bars indicate standard error of the mean, computed in a circular fashion. **(B** and **E)** Phase Response Curve, showing start phase vs phase change from predicted phase (in radians) for Fz in experiment 3 (**B**) and Pz in experiment 4 (**E**). Horizontal line indicates zero. Vertical coloured lines indicate where actual conditions fell in their stimulus onset phase. Error bars indicate standard error of the mean, computed in a circular fashion. **(C** and **F)** Group phase and resultant for each of the 10 starting phase bins. Black line indicates resultant of all phase bins. Coloured lines indicate trajectory of each phase bin, across the preceding 50 ms, for Fz in experiment 3 (**C**) and Pz in experiment 4 (**F**). All figures are shown for delays 20 ms, 100 ms, 150 ms, and 200 ms.

### Study 3. Application to Sleep Onset – Functional Implications

#### αCLAS during the awake-to-sleep transition modulated alpha frequency in a phase-dependent manner

The phase-dependent effects observed in the previous studied suggest that phase may be leveraged using *αCLAS* to modulate explicit brain processes that rely on dynamics associated with alpha oscillations. As the brain transitions away from wakefulness during the process of falling asleep, alpha oscillations undergo a dramatic disappearance across the cortex. This process first systematically described in the 1930’s[47,48] is still employed to define the first stage of non-rapid-eye-movement (NREM) sleep. If our *αCLAS* modulation of the alpha rhythm is functionally meaningful, then it might also modulate alpha-associated processes in a phase-dependent way.

To investigate this, sixteen participants took part in a within-subjects, sham-controlled, cross-over nap study (Study 3), in which repetitive-*αCLAS* was once more employed, phase-locked to alpha at Fz, where we previously demonstrated an effect of *αCLAS*. In separate visits, participants were exposed to three conditions (sham, i.e. no sound, pre-peak and pre-trough of alpha), and given a 31-minute opportunity to sleep, the first minute served as baseline, after which sounds played for 15 minutes (in the stimulation conditions), followed by 15 minutes of silence (**Figure 1B**). This design allowed us to explore online (i.e. during) and offline (i.e. post-stimulation) effects of phase-locked alpha stimulation on metrics of sleep and brain dynamics.

We observed a striking and sustained difference in alpha frequency at the phase-locking electrode (i.e. Fz) between conditions during stimulation (i.e. initial 15 min), which ceased when stimulation ended (**Figure 7Ai**). The increase in alpha frequency during pre-peak compared to pre-trough condition was observed for 15 out of 16 participants (**Figure 7Aii**) and both conditions were significantly different in frequency to sham. Once stimulation ceased, so did the differences in frequency (**Figure 7Aiii**). Overall, the direction of these differences was consistent with Study 1, with pre-trough slowing and pre-peak accelerating the alpha frequency.

**Figure 7.**
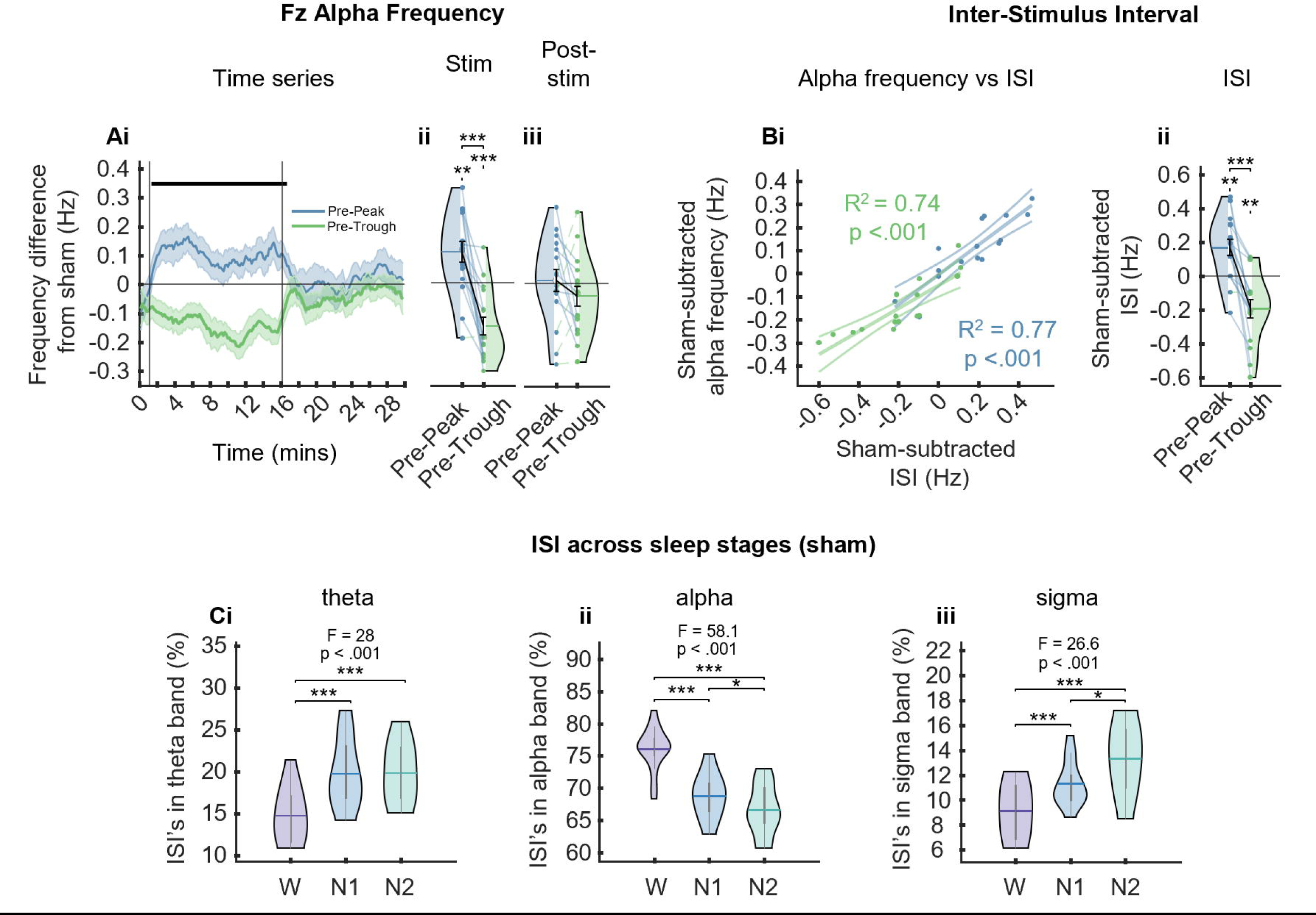
Alpha Frequency and Inter-Stimulus Interval during Nap study (Study 3) **(A)** Sham-subtracted Fz alpha frequency **(i)** across time, error bars are SEM vertical lines represent stimulation start and stop; thick horizontal bar indicates timepoints at which *p* ≤ 0.5, as calculated by t-test **(ii)** collapsed across the stimulation period; **(iii)** collapsed across the post-stimulation period. **(B) (i)** Sham-subtracted ISI vs sham-subtracted alpha frequency. Lines and statistics derived from simple linear regression for each condition; **(ii)** average sham-subtracted ISI **(C)** percentage of sham ISI’s in different sleep stages falling in the **(i)** theta band (4-7 Hz); **(ii)** alpha band (8-12 Hz); and **(iii)** sigma band (12-16 Hz) for the sham condition (see **Figure S10** for all conditions). In-figure text shows p values and F values from linear mixed effects models output. *** *p* < 0.001, ** *p* < 0.01, * *p* < 0.05, t-tests.

As might be expected, the ISI between pulses of sounds showed a strong linear relationship with alpha frequency (**Figure 7Bi**). In a truly closed-loop experiment [25], it can become difficult to distinguish the direction of causality – did shorter ISI’s speed up the brain’s rhythms or did faster brain rhythms lead to shorter ISI’s? Using the data from the sham condition, in which sound triggers were locked to pre-peak, but no sound was played, allowed us to disentangle this conundrum. These dummy sound pulses were locked to the same phase as the pre-peak condition (**Figure S10**), but the ISIs differed, indicating that the ISI was indeed dependent on the brain’s responses to sound, as opposed to the particular phase targeted. Furthermore, the distribution of ISIs differed between vigilance states (**Figure 7C**, **Figure S11**). During sleep there was an increase of ISIs in the theta and sigma range, representative of the theta oscillations and spindles that characterise shallow sleep (N1/N2). Whilst a greater proportion of alpha band ISI’s was seen during wakefulness, further suggesting that the ISI is dependent on the brain’s physiology.

#### αCLAS impacts sleep *macro*structure in an alpha phase-dependent manner

Sleep scoring showed the expected trajectory during the nap; all participants began in an awake state and the majority were asleep (defined by any stage of NREM) after approximately 10 minutes (**Figure 8Ai**). We observed some awakenings once stimulation ceased (dotted line in the timeseries plots in **Figure 8**) with some participants reported being asleep until the sounds stopped. Although unintended, this is consistent with the finding that the sleeping brain remains sensitive to changes in environmental stimuli [49]. Aside from these awakenings, all three conditions showed a comparable trajectory and total sleep time (TST) did not significantly differ between conditions during either the stimulation or post-stimulation periods (**Figure 8A**).

**Figure 8.**
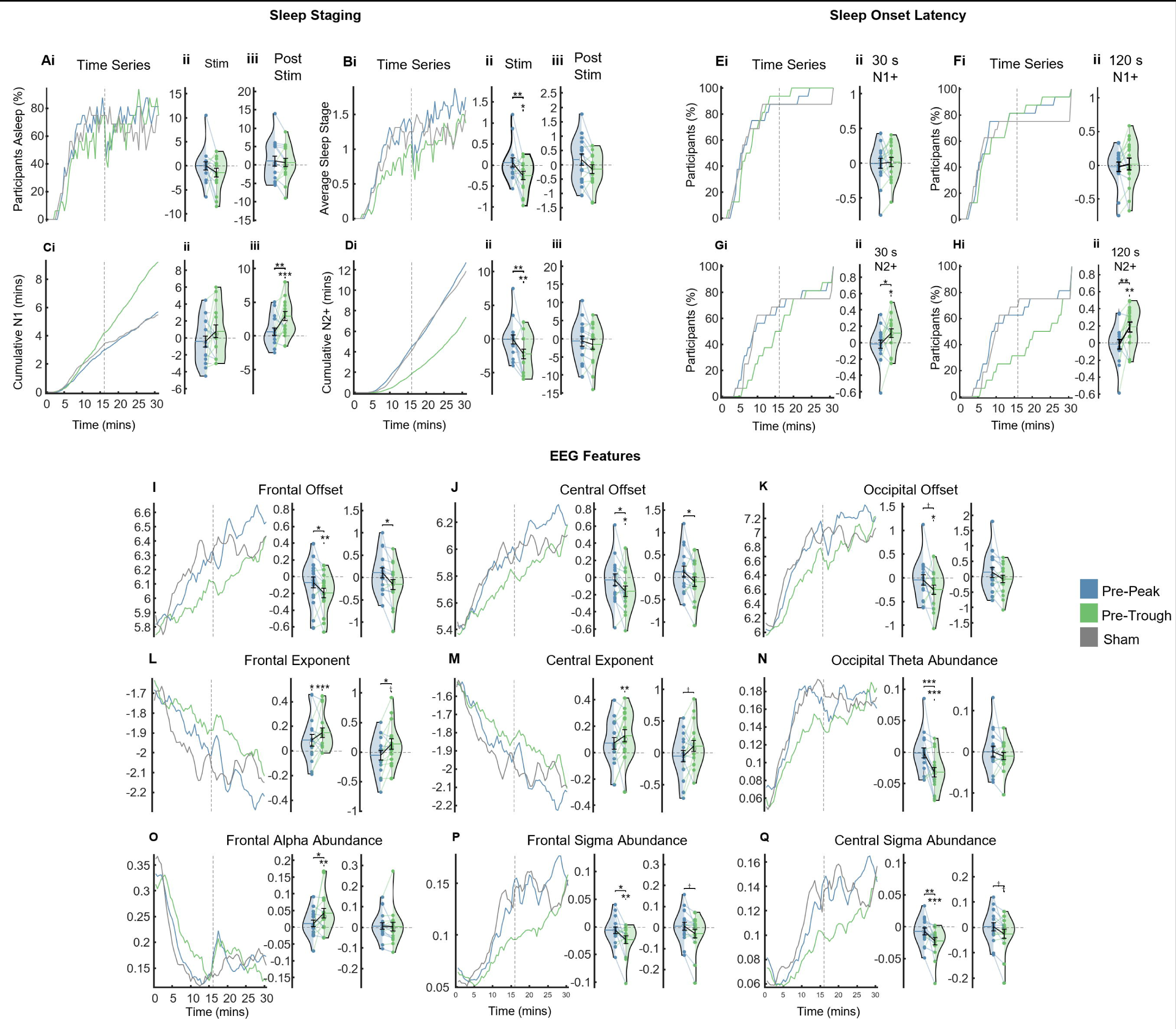
Sleep Structure and EEG features. **(A) (i)** Timeseries of the percentage of participants in any stage of vigilance (each state was given a numerical value: awake - 0, N1/NREM1 – 1, N2/NREM2 - 2, N3/NREM3 – 3, and no REM sleep was observed). Dashed line indicates end of stimulation **(ii)** Sham-subtracted total sleep time (TST) collapsed across stimulation period **(iii)** Sham-subtracted TST collapsed across post-stimulation period. **(B) (i)** Average sleep stage timeseries. Dashed line indicates end of stimulation **(ii)** Sham-subtracted average sleep stage collapsed across stimulation period **(iii)** Sham-subtracted average sleep stage collapsed across post-stimulation period. **(C) (i)** Cumulative N1 sleep timeseries. Dashed line indicates end of stimulation **(ii)** Sham-subtracted time in N1 sleep collapsed across stimulation period **(iii)** Sham-subtracted time in N1 sleep collapsed across post-stimulation period. **(D) (i)** Cumulative N2+ (i.e. N2/N3) sleep timeseries. Dashed line indicates end of stimulation **(ii)** Sham-subtracted time in N2+ sleep collapsed across stimulation period **(iii)** Sham-subtracted time in N2+ sleep collapsed across post-stimulation period. **(E) (i)** Time series of participants (%) having had at least 30 seconds of sleep **(ii)** Sham-subtracted log-transformed latency of sleep onset (any stage, at least 30 seconds). **(F) (i)** Time series of participants (%) having had at least 120 persistent seconds of sleep **(ii)** Sham-subtracted log-transformed latency to the onset of the first persistent 120 seconds of sleep (any stage). **(G) (i)** Time series of participants (%) having had at least 30 seconds of N2+ sleep **(ii)** Sham-subtracted log-transformed latency of N2+ sleep onset. **(H) (i)** Time series of participants (%) having had at least 120 persistent seconds of N2+ sleep **(ii)** Sham-subtracted log-transformed latency to the onset of the first persistent 120 seconds of N2+ sleep. **(I – Q)** Timeseries of EEG features (left) averaged across participants for each condition that showed a significant main effect of phase condition. Sham-subtracted features collapsed across stimulation period (middle) and post-stimulation period (right). Note: statistics were run prior to sham subtraction. Post-hoc comparisons were only carried out when a main effect of condition was seen in LME. Significance bars between violins indicate a difference between stimulation conditions, significance marks over violins indicate a difference from sham. *** *p* < 0.001, ** *p* < 0.01, * *p* < 0.05, † *p* < 0.1, t-tests. Each violin shows a dot per participant, per condition, horizontal lines indicate the mean.

Average sleep stage increased over time, and was significantly different between conditions – pre-trough stimulation showed a diminished sleep depth, but only during stimulation (**Figure 8B**). When broken down further into N1 and N2+ (epochs labelled as N2/N3) these differences became more apparent – the trajectory of cumulative N1 sleep clearly trailed off in pre-peak and sham conditions, as participants transitioned to deeper stages of sleep, whilst in pre-trough the amount of N1 continued to accumulate, with significantly more N1 sleep seen post-stimulation (**Figure 8C**). Similarly, a diminished trajectory of cumulative N2+ sleep was seen in the pre-trough condition, amounting to significantly less N2+ sleep during the stimulation period when compared to pre-peak and sham (**Figure 8D**).

Latency to onset of the first epoch of sleep (any stage) did not differ between conditions, nor did latency to onset of the first persistent 2 minutes of sleep (**Figure 8E**, **Figure 8F**). However, participants took significantly longer to reach the first epoch of N2+ in pre-trough stimulation (**Figure 8G**) and this effect was greater still when looking at latency to the first persistent 2 minutes of N2+ sleep (**Figure 8H**). These results show that pre-through stimulation impacted sleep macrostructure, relative to sham, while pre-peak stimulation did not.

#### αCLAS impacts sleep *micro*structure in an alpha phase-dependent manner

Sleep scoring is a useful way to broadly classify sleep physiology, but its drawbacks are well known. Namely, the brain probably exists on something of a continuum rather than conforming to four or five discrete states. To arrive at a finer grain measure of EEG we characterised both rhythmic, i.e. oscillatory, and arrhythmic, i.e. aperiodic or 1/f, EEG activity using eBOSC [50].

Several eBOSC features showed a significant main effect of *αCLAS* phase condition (**Figures 8I-8Q** display their trajectories, **Supporting Tables 1** and **2** for full statistics). These features captured many of the known dynamics of sleep onset – steepening of power spectra (offset of 1/f slope increases, exponent decreases), accompanied by a sharp decrease in alpha oscillations and the appearance of theta oscillations and sleep spindles (**Figure S12** for sleep-stage comparison and **Figure S13** for trajectory across time of all EEG features).

The majority of these metrics confirmed findings from the sleep scoring –during pre-trough *αCLAS* features which increase in magnitude during the natural progression of sleep (offset in all regions, occipital theta abundance, frontal and central sigma abundance) were reduced, whilst features which decrease with sleep were greater (frontal and central exponent, and frontal alpha abundance). Especially compelling were the differences in frontal and central sigma abundance, representative of spindles – a well-established hallmark of sleep. Frontal exponent was significantly shallower than sham during both types of stimulation, but this was the only metric by which sham and pre-peak *αCLAS* could be distinguished.

In sum, the trajectory of sleep was modified by *αCLAS* in an alpha phase-dependent manner. Specifically, whilst the onset and duration of sleep did not differ between conditions, its composition did; a ‘shallower’ sleep was observed when sounds were repeatedly administered just prior to the troughs of alpha oscillations, relative to pre-peak locked sounds and the absence of sounds.

## DISCUSSION

Across three independent studies we have shown: (i) *αCLAS* can induce phase-dependent modulations of alpha activity in a spatially specific manner (Study 1), (ii) this effect is dependent on oscillation amplitude (Study 2), (iii) sounds delivered at different phases influence the trajectory of alpha oscillations (Study 2), and (iv) the transition to sleep can be modulated by *αCLAS* (Study 3).

We first demonstrated that phase-dependent responses to auditory stimulation of the alpha oscillation can be elicited in the resting brain. By employing hd-EEG and targeting two distal cortical locations, we observed that repetitive-*αCLAS* modulated the alpha rhythm in a spatially specific manner, with changes to alpha power, frequency, and connectivity present when targeting the frontal, but not the parietal location. Whilst the frequency changes may appear very small they are consistent with, or greater than, those derived from the same analysis method elsewhere in the literature [7,35,51].

Previous studies on phase-dependencies of neural oscillations consider phase an index of ‘cortical excitability’, i.e. a cyclical measure of the cortex’s sensitivity to stimuli. These studies focused primarily on secondary, or higher order, outcomes (perception, for example [5,7,8,52–55]) of stimuli delivered at a particular phase without making explicit predictions on the oscillation’s response, or they expected a straightforward change in the amplitude of the targeted oscillation [56], akin to observations from sound stimulation to the slow oscillations of sleep [27–29]. Whilst our stimulation effects agreed with the literature, insofar as they are phase-dependent, we found they could be better explained by changes to the *frequency* of oscillations, rather than amplitude.

Striking phase-dependent changes in alpha-band connectivity were also observed, which we suggest are at least partially consequent of this localised frequency change, but may also have implications for theories of intra-brain information transfer [39,40].

By targeting the alpha rhythm at multiple phases while delivering single-pulse *αCLAS* we tested whether a phase-reset mechanism would provide a parsimonious explanation to the observed alpha frequency modulation. Our results support this hypothesis and were consistent with an oscillator model, in which a perturbation will advance or delay an oscillation to varying degrees depending on the phase at which it is administered [41,45,46]. While there is no unanimous consensus on how related or unrelated evoked potentials are to phase reset of oscillations [57], our results cannot simply be explained by the response evoked by the auditory stimulus, as we observed similar auditory evoked potentials across phase conditions. In addition, our results showed that the frequency dependency on phase was maintained, for short (Study 1) and sustained periods of stimulation (Study 3).

Despite, phase-locking with high spatial selectivity, modulation of alpha activity was limited to phases targeted at the frontal region. We put forward two complementary explanations for this phenomenon supported by our findings. The first lies in the endogenous regional differences in alpha amplitude, i.e. higher in parietal than frontal locations. Indeed, our results demonstrate that a parietal phase-reset could only be observed in instances of low alpha power, and that alpha power and phase reset magnitude are strongly related. This is also in line with oscillator theory, which proposes a phase reset is dependent on the magnitude of both the oscillator and the perturbation, and agrees with prominent theories that the phasic inhibition exerted by alpha oscillations is proportional to their amplitude [15,58]. Secondly, the differential in the magnitude of AEPs at frontal and parietal regions could be a contributory factor to regional differences. Future studies employing visual stimuli phase-locked to parietal alpha oscillations could help disentangle the contribution of evoked potential magnitude and oscillation amplitude in phase-dependent responses, since visual evoked potentials are larger over the visual cortex in spite of higher amplitude oscillations.

Finally, in addition to the immediate electrophysiological consequence of *αCLAS,* we have shown the functional significance of this approach, during the transition to sleep. Auditory stimuli modulated alpha frequency, once again, and sleep depth on a phase-dependent basis. We, once more, put forward two hypotheses. In Study 1, in addition to a frequency change, we showed that pre-peak *αCLAS* predominantly caused a decrease in alpha power, whilst pre-trough *αCLAS* did the opposite, i.e. an increase in alpha power. Accordingly, during the transitionary period of N1 sleep, pre-trough *αCLAS* may have impeded the disappearance of alpha, preventing sleep from progressing further, whilst pre-peak *αCLAS* did not. Alternatively, differences in sleep depth may simply reflect phasic differences in sensitivity to sound. In this scenario, pre-trough locked sounds arrived at a more suitable phase at which to perturb the brain and in consequence the sleep process, in a similar vein to the cyclic excitability put forward in the visual neuroscience literature [5,6,8,14,15,52,53,59]. Studies using intracranial electroencephalographic (iEEG) recordings in humans or animal models could help to elucidate the direct neural response and offer further mechanistic insights.

A general limitation of our studies is that our closed-loop stimulation did not account for variations in the endogenous alpha power. This was mitigated by the fact that participants had their eyes closed, a state of abundant alpha oscillations, and variations in alpha power were used to our advantage in Study 2. However, in Study 3, we see that ISIs deviate to other frequencies when alpha oscillations are reduced as participants fall asleep. While we cannot completely rule out an effect of ISI, or stimulation frequency in Study 3, such that the impact of sound on sleep is only phase-dependent in the sense that the frequency of sound was phase-dependent, the differences in ISI (0.4 Hz on average) are very slight, rendering this explanation unlikely. Future studies can implement closed-loop strategies that include a detection threshold, thus limiting stimulation to periods when the target rhythm is most prominent. Another helpful addition may be the inclusion of a random-sound condition – in Study 3 we were unable to say for certain whether the pre-peak represented a moment of decreased sensitivity to sound, or pre-trough a window of increased sensitivity. Finally, we used a mastoid reference for real-time phase-locking. The use of an on-line Laplacian reference, as seen in some other closed-loop experiments [21–23], may afford stronger, and further localised effects.

Alpha oscillations are involved in a number of fundamental processes. Here we demonstrate that sound can be used to effect phase-dependent modulation of their activity. Future research could extend our findings into potential clinical applications. For example, by countering the decline in alpha frequency observed in older age and cognitive decline/dementia [60,61] or facilitating the increase in alpha frequency observed during increases in cognitive load [62]. There remains much to be explored, regarding the application of *αCLAS* to neural oscillation-dependent behaviours, but we propose that sound can provide more utility in this context than previously considered, particularly if stimulation is used to address the brain on its own terms, through closed-loop approaches.

## MATERIALS AND METHODS

### Participants

#### Study 1

58 participants took part in this study, 25 in experiment 1 and 31 in experiment 2. Four participants were excluded from experiment 1 due to missing triggers, and 1 participant was removed due to poor quality EEG. Three participants were excluded from experiment 2 due to missing triggers. This left a sample of 20 participants in experiment 1 (14 female, mean ± std age = 22.55 ± 3.81 years) and 28 participants in experiment 2 (18 female, mean ± std age = 20.77 ± 2.49 years). Five participants overlapped between the two experiments.

#### Study 2

15 participants took part in this study, 8 in experiment 3 (7 female, mean ± std age = 24.50 ± 5.90 years) and 7 in experiment 4 (6 female, mean ± std age = 26.71 ± 5.82 years). Five participants took part in both experiments. All participants were included in the analysis.

#### Study 3

25 participants took part in this study, 9 of whom were excluded (1 for sleep-onset REM, 2 for technical issues, 5 for missing visits due to illness or other reasons, 1 for night shift between visits). The final sample was composed of 16 participants (14 female, mean ± std age = 22.38 ± 2.74 years).

Across the three studies, all participants self-reported no history of neurological or psychiatric illness, normal hearing, and a body mass index of less than 30. Participants taking part in Study 3 were asked not to consume caffeinated drinks on the morning of the experiment and refrain from drinking alcohol the nights before the experiment.

Participants gave written informed consent. The study conforms to the Declaration of Helsinki and received ethical approval from the University of Surrey’s ethics committee.

### Experimental Design

#### Study 1

Participants sat in a comfortable armchair in a soundproof room with constant LED light (approx. 800 Lux). Participants were exposed to 8 runs (2 runs per phase) of auditory stimulation phase-locked to either Fz (experiment 1) or Pz (experiment 2). Each run comprised 5 blocks of 30 s stimulation, interleaved with 10 s of silence, and started with 10 s of silence. The order of the runs was counterbalanced across participants using a balanced Latin square. Each run lasted 200 s. At the beginning of each run participants were asked to close their eyes and relax. At the end of each run participants were told to open their eyes and electrode impedance was checked and, if necessary, adjusted. This procedure helped to avoid participants falling asleep during the session. In total, each session lasted approximately 2 h.

#### Study 2

The study was conducted under the same laboratory conditions as Study 1. However, instead of blocks of continuous sound stimulation, participants were exposed to isolated bursts of sound. Participants were exposed to 8 runs (2 per phase targeted) of auditory stimulation phase-locked to either Fz (experiment 3) or Pz (experiment 4). Each run comprised 96 pulses of pink-noise, interleaved with between 1.5 and 5.5 s of silence (randomly generated, uniformly distributed, mean ± std = 3.5 ± 0.88 s). The order of the runs was counterbalanced across participants using a balanced Latin square. Each run lasted around 350 s. Total number of pulses per condition was 192, and the total duration of the session was around 2 h.

#### Study 3

Participants attended the Surrey Sleep Research Centre lab for a total of four sessions, the first of which was an adaptation visit, to allow participants to acclimatise to the sleep centre and protocol. Visits were separated by at least 2 days. Each session had an approximate duration of 3 h and in addition to the nap included a memory task, Karolinska Drowsiness Test (KDT) and Karolinska Sleepiness Scale (KSS), which were not analysed here. Throughout the study, starting at the adaptation visit, participants were requested to use a triaxial GENEActiv accelerometer (Activinsights, Kimbolton, UK), on their non-dominant wrist, and complete sleep diaries, which were checked to confirm a regular sleep schedule. Participants also completed a Pittsburgh Sleep Quality Index (PSQI) questionnaire to assess sleep health.

During the nap portion of the visit, participants lay supine in bed in a temperature-controlled, sound-attenuated, pitch-dark, windowless room for the duration of the nap opportunity (i.e. 31 min). Participants were asked to keep their eyes closed for the entire nap period, regardless of whether they were asleep or not. All visits took place in the morning, starting at 07:00, 08:00 or 09:00 h, with the naps starting approximately 2 h into the visit. The time of each visit, and hence each nap, was consistent across visits for each participant. Participants were blinded to the condition, as was the researcher providing instructions to the participant.

### Phase-Locked Auditory Stimulation

Real-time closed-loop phase-locked auditory stimulation was delivered using a custom-made device as described in [34]. This device utilised three electrodes; a signal electrode (placed at Fz or Pz, depending on the experiment), a reference and ground (both placed at the right mastoid). Shielded electrodes (8mm, BIOPAC Systems Inc.) were used for the signal and reference, and an unshielded electrode was used for the ground (8mm, BIOPAC Systems Inc.). The signal was filtered using a 7.5-12.5 Hz bandpass, and the phase of the resultant signal was computed in real-time using the endpoint-correct Hilbert Transform (ecHT). The device used for Studies 1 and 2 operated at a sampling rate of 500 Hz, whilst that used in Study 3 operated at 826 Hz. Sounds were 80 dB, 20 ms pulses of pink noise, delivered to the participants via earphones (Etymotic model ER-2 in Studies 1 and 2 and Shure model SE215-CL-EFS in Study 3). Sounds were phase-locked to 4 phases of alpha in Studies 1 and 2 (60°, 150°, 240° and 330°) and two phases in Study 3 (150° and 330°). In the sham condition in Study 3, phase-locking was still computed, and markers were logged for all ‘stimuli’ (dummy targeting 330°), but the volume was set to 0 dB ensuring that no sounds were played to the participant.

### EEG data acquisition and sleep monitoring

#### Studies 1 and 2

High-density (hd)-EEG recordings were acquired in parallel with the phase-locking EEG device. Recordings were obtained using an actiCHamp-Plus amplifier (Brain Products GmbH) and 128-channel water-based R-NET electrode cap (EasyCap GmbH, Brain Products GmbH) positioned according to the international 10-20 positioning system. Electrode impedances were below 50 kΩ at the start of each recording, as per the manufacturer’s guideline, and were adjusted between each run, to ensure accurate impedances throughout the recording. All channels were recorded referenced to Cz, at a sampling rate of 500 Hz.

#### Study 3

Sleep recording equipment was acquired in parallel with the phase-locking EEG device. Sleep recording equipment was a Somnomedics HD system with Domino software (v2.8), sampled at 256 Hz (S-Med, Redditch, UK) using the American Academy of Sleep Medicine (AASM) standard adult EEG montage. The system included 9-channel EEG placed according to the 10-20 system at F3, F4, C3, C4, O1, O2, left mastoid (A1) and right mastoid (A2), and referenced to Cz, as well as submental Electromyography (EMG), electro-oculogram (EOG), and electrocardiogram (ECG).

The ecHt device was programmed to send triggers at the beginning of stimulation blocks in Study 1, which were recorded in the BrainVision Recorder software (Brain Products GmbH) alongside EEG data, via a TriggerBox (Brain Products GmbH). In addition, in Studies 1 and 2, we used a StimTrak (Brain Products GmbH) that logged a trigger whenever a sound was played, allowing us to assess phase-locking in the hd-EEG data and to time-locked evoked potentials. In Study 3, an analogue marker was sent from the ecHT device to the S-Med PSG system, so as to align EEG recordings according to the start of the nap.

### EEG Pre-processing

All EEG data were pre-processed and analysed in MATLAB 2021a (Mathworks, Natick, MA). EEG filtering was carried out using the FieldTrip toolbox [63], all other pre-processing was carried out using EEGLAB v2022.0 [64]. All topoplots and related statistics were produced using FieldTrip [63].

Hd-EEG data were high-pass filtered (1 Hz) and notch filtered (50 Hz, 100 Hz), before automatic subspace reconstruction (ASR) was used to remove noisy channels and artefacts. This was followed by independent component analysis, to identify and remove components related to ocular and cardiac artefacts, in Study 1. EEG data from Study 3 were filtered between 1 and 30 Hz, before artefacts were manually identified and removed. The hd-EEG data was subjected to a scalp current density re-referencing, whilst the EEG data from study 3 remained referenced to Cz.

### EEG Analysis

Accuracy of phase-locking was calculated for each channel, participant, condition and experiment using the following method. For every stimulus, the EEG was epoched in 1 s windows up to and including stimulus onset, before the ecHT algorithm was applied, and the final phase estimate in each window was taken. The ecHT algorithm minimises the Gibbs phenomenon by applying a causal bandpass filter to the frequency domain of an analytic signal, thus selectively removing the distortions to the end part of the signal. This strategy has been shown to provide accurate computations of the instantaneous phase and envelope amplitude of oscillatory signals [34]. The resultant vector length of these values was then computed, giving a value between 0 and 1, representing perfectly uniform and perfectly unimodal circular distributions respectively. The accuracy of phase-locking for the group was summarised by computing the average resultant of all participants, and computing a z-test. Z-test was calculated using phase angle and resultant per participant, and the respective target phase angle (i.e. 60°, 150°, 240°, 330° depending on the condition).

For all analyses regarding power, the same method was used; power was computed in 1-second epochs, 2-30 Hz, in steps of 0.1 Hz, Hamming window, using the ‘pwelch.m’ function. For all analyses regarding frequency, Cohen’s frequency sliding method was used [35].

#### Study 1

Frontal and parietal regions of interest (ROI) were defined *a priori*, and included four electrodes, the phase-locking electrode and the three surrounding electrodes. The frontal cluster comprised AFz, Fz, AFF1h, and AFF2h. Whist the parietal ROI consisted of Pz, POz, PPO1h, and PPO2h.

For all Study 1 analyses of power spectra, percentage changes from ‘off’ to ‘on’ were computed for every frequency bin. Before running statistics and plotting time-frequency representations, data were smoothed *across time*, using a moving mean window of 4 seconds. ‘movmean.m’ function, with a window argument of ‘[0 4]’, meaning each second in the plot was an average of that second and the 4 seconds that followed.

Statistics were computed for these time-frequency representations using either one-way ANOVA or t-tests as appropriate. Statistically significant (*p* < 0.05) contiguous clusters of 10 or more time-frequency points were kept and considered significant, and smaller clusters were discarded to control for multiple comparisons. Individual alpha frequency (IAF) was calculated at the respective phase-locking electrode (i.e. Fz/Pz) using the frequency sliding method [35].

Connectivity was computed for four frequency bands (delta 1-4 Hz, theta 4-7 Hz, alpha 8-12 Hz, beta 13-30 Hz), for every second of clean EEG data. Two metrics were used, phase-locking value (PLV) and phase lag index (PLI). In both cases, the following was carried out for each channel pair: the data from each channel was filtered in the band of interest using a 2^nd^ order Butterworth filter; phase difference between the two channels was estimated for all time points using Hilbert transform derived phase estimates; for each second, the resultant vector length of phase differences was computed. The only difference between PLV and PLI was that in the case of PLI, phase differences of 0 mod π were discarded prior to computing the resultant. When comparing conditions, these measures of connectivity were averaged across the ‘on’ (sound stimulation) period.

When computing stimulation-induced changes to connectivity, these measures were separately averaged across the ‘on’ and ‘off’ periods before comparing the two.

#### Study 2

Broadband auditory evoked potentials were computed by filtering the data between 1 and 40 Hz (2^nd^ order Butterworth), and averaging over all trials, per condition, per participant. Narrowband auditory evoked potentials (both amplitude and phase) were computed by applying the ecHT algorithm – a 1 s window was slid in increments of 1 data point (2 ms), each window was subjected to ecHT, and the estimate for the final timepoint in that window was assigned to that timepoint in the data.

To assess unimodality of phase angles between conditions (and hence, a phase reset), a Rayleigh test was carried out for each timepoint, using the average phase angle and resultant for each participant, for each condition. Timepoints at which the Rayleigh p≤0.05 were considered to have significant unimodality.

Trials were sorted into octiles on the basis of pre-stimulus (-200 to -100 ms) alpha power, (mean ± SD number of trials per octile per participant: experiment 3: 22.69 ± 0.998; experiment 4: 21.86 ± 2.37). This allowed us to assess the effect of oscillation amplitude on phase-reset.

The aforementioned phase estimates were also used to sort all evoked potentials by stimulus onset phase, into ten 36° bins centred on 0°, 36°, 72°, 108°, 144°, 180°, 216°, 252°, 288°, and 324° (mean ± SD number of trials per phase bin per participant; experiment 3: 74.02 ± 1.45, minimum: 67.1, maximum: 80; experiment 4: 72.19 ± 2.72, minimum: 67.1, maximum: 80.4). This allowed phase response curves to be computed at various latencies, by plotting starting phase bin against end phase, and to derive phase transfer curves by plotting starting phase bin against change from expected phase (assuming a 10 Hz oscillation).

#### Study 3

Sleep was scored by an experienced sleep technician blinded to the experimental condition, according to AASM guidelines. In order to better assess the dynamics of the ecHT’s stimulation, we computed inter-stimulus intervals ISI’s by taking the difference in time between each stimulus and the stimulus that proceeded it, before converting this value to Hz. ISI’s greater than 0.5 s (<2 Hz) were discarded before the average ISI was computed by taking the median. The proportion of ISI’s which fell in canonical bands (theta 4-7 Hz, alpha 8-12 Hz, sigma 12-18 Hz) was then computed, to demonstrate that the behaviour of the stimulator followed the oscillatory features of the sleep stages.

We employed eBOSC [50] to estimate both rhythmic, i.e. oscillatory, and arrhythmic, i.e. aperiodic or 1/f, EEG activity. For eBOSC analysis, for each channel, EEG data were divided into 30 s epochs, since, during a dynamic process such as sleep onset, a single fit and threshold for oscillation detection is unlikely to be optimal. For each of these epochs, eBOSC was run between 1-30 Hz, in steps of 0.25 Hz, excluded frequencies for background 1/f fit 7:16 Hz. Only oscillations with a duration of at least 3 cycles were analysed. The abundance computed was then averaged within canonical bands for each epoch. This was done for all channels (F3, F4, C3, C4, O1, O2) before further averaging into regions (frontal: F3, F4; central: C3, C4; occipital: O1, O2).

### Statistical Analyses

Unless stated otherwise, statistics were run in the following manner. Cluster-corrected permutation t-tests or ANOVA were run on FieldTrip [63] and used for topographical representations. Linear mixed-effects models (LMEM), in which participant was a random effect, were used throughout (for all but one violin plot – Figure 4B). Where these LMEM registered a significant main effect (*p* ≤ 0.05) post-hoc comparisons between all groups were carried out using the ‘emmeans’ toolbox (https://github.com/jackatta/estimated-marginal-means), so as to control for multiple comparisons. Study 3 features a number of ‘stim minus sham’ plots – in these cases, an LMEM was run on the data from the three groups, prior to subtracting sham, and the sham-subtraction was applied for visualisation purposes. All circular statistics were conducted using the ‘circstat’ toolbox [65].

## Code and Data Availability

Data and key scripts will be made available on Gitlab.

## Author contributions

H.H., N.G., D.-J.D. and I.R.V. designed the experiments. H.H., B.L. and R.D. performed the experiments and recorded the data. H.H. curated the data and performed the data analyses. E.R. and V.J. assisted with formal data analysis. I.R.V. and D.-J.D. supervised the investigation. N.G. developed the methodology. H.H. and I.R.V wrote the initial draft. All authors reviewed and edited the manuscript.

## Acknowledgments

We thank Sarah Leslie and Adrian Cheung for their contributions to data collection, Giuseppe Atzori for scoring the sleep EEGs, David Wang for his technical support, and Ullrich Bartsch for fruitful discussion of our results.

H.H. is supported by the University of Surrey’s Doctoral College Scholarship Award.

I.R.V. is supported by the Biotechnology and Biological Sciences Research Council (BB/S008314/1). D.-J.D is supported by the UK Dementia Research Institute which receives its funding from DRI Ltd, funded by the UK Medical Research Council, Alzheimer’s Society and Alzheimer’s Research UK. N.G. was funded by the UK Dementia Research Institute (UK DRI)—an initiative funded by the Medical Research Council, Alzheimer’s Society and Alzheimer’s Research UK, Wellcome Trust fellowship (097443/Z/11/Z), Science & PINS Award for Neuromodulation, and NIHR IBRC Confident in Concept Award.

## Supporting Information Legends

**Figure S1. Phase-locking Accuracy Across Conditions**

Topography of phase-locking accuracy (average resultant) across 4 conditions of two experiments. White marks indicate channels at which resultant >0.5 and p <0.05. Black marks indicate channels at which resultant <0.5 and p <0.05. p values from Bonferroni-corrected z-test for non-uniformity.

**Figure S2. Power change ANOVA for experiment 1**

Topography of permutation ANOVA stats for each 0.5 Hz bin. White dots show significant main effect of phase-targeted, cluster-corrected *p* <0.05

**Figure S3. Alpha band power changes experiment 1**

**(A)** topography of permutation ANOVA stats for each .5 Hz bin in experiment 1. **(B)** pre-peak vs pre-trough, positive values indicate pre-trough is greater **(C)** post-peak vs post-trough, positive values indicate post-peak is greater. White dots show significant main effect of phase targeted, cluster-corrected p <0.05.

**Figure S4. Frequency estimates form frontal ROI in experiment 1**

For all plots, mixed effects models were run: [frequency/ISI ∼ condition + (1|Participant)]. Post-hoc t-tests were run for those which showed a statistically significant effect of condition (*p* < 0.05). * *p*<0.05, \*\**p*<0.01, \*\*\**p*<.001.

**Figure S5 Power change ANOVA for experiment 2**

Topography of permutation ANOVA stats for each 0.5 Hz bin. White dots show significant main effect of phase-targeted, cluster-corrected *p* <0.05

**Figure S6. Alpha band power changes experiment 2**

**(A)** topography of permutation ANOVA stats for each .5 Hz bin in experiment 1. **(B)** pre-peak vs pre-trough, positive values indicate pre-trough is greater **(C)** post-peak vs post-trough, positive values indicate post-peak is greater. White dots show significant main effect of phase targeted, cluster-corrected p <0.05.

**Figure S7. Frequency estimates form parietall ROI in experiment 2**

For all plots, mixed effects models were run: [frequency/ISI ∼ condition + (1|Participant)]. Post-hoc t-tests were run for those which showed a statistically significant effect of condition (*p* < 0.05). * *p*<0.05, \*\**p*<0.01, \*\*\**p*<.001.

**Figure S8. Topography of permutation ANOVA stats for each frequency band in experiment 1.** White dots show significant main effect of phase-targeted, cluster-corrected p <0.05.

**Figure S9. Topography of permutation ANOVA stats for each frequency band in experiment 2.** White dots show significant main effect of phase-targeted, cluster-corrected p <0.05.

**Figure S10. Phase accuracy plots for the ecHT electrode in the three conditions in Study**. **3.** Each line represents a participant, length of line indicates resultant (between 0 and 1). Phase accuracy is high in all three conditions. Phases are the same for pre-peak and sham. During sham, markers were recorded for each sound stimulus, but the volume was zero.

**Figure S11. Inter stimulus intervals per sleep stage.** Violin plots show the percentage of ISI’s in each frequency band, in each sleep stage, in each condition. Stats indicate output of linear mixed effects model [ISI_percentage ∼ sleep_stage + (1|participant)]. * p <0.05, ** p<0.01, ***p<0.001.

**Figure S12. eBOSC features per sleep stage for the sham condition**

**(A)** The offset of the aperiodic component. **(B)** The exponent of the aperiodic component. **(C)** The abundance of theta oscillations. **(D)** the abundance of alpha oscillations. **(E)** The abundance of sigma oscillations. Averages are shown for each feature, each region, each participant, for the sham condition.

Stats indicate output of linear mixed effects model [eBOSC_feature ∼ sleep_stage + (1|participant)].

Bars indicate results of t-tests, in cases of a significant main effect, three contrasts were run: N1-W, N2-N1,N3-N2 * p <0.05, ** p<0.01, ***p<0.001.

**Figure S13. Time series for each eBOSC features.** Average time series are shown for each feature, each region, for each condition. Data was smoothed using a 2-minute moving mean window

**Table S1** Results of linear mixed-effects models of the form: [eBOSC_feature ∼ condition + (1|participant)] for the *stimulation period*. Where a main effect of condition was found (*p* <0.05), post-hoc contrasts were carried out between estimated means of model. Df, degrees of freedom; Df.res, Residual Degrees of Freedom; F, F-test statistic; p, p-value; x, not computed.

**Table S2** Results of linear mixed-effects models of the form: [eBOSC_feature ∼ condition + (1|participant)] for the *post-stimulation period*. Where a main effect of condition was found (*p* <0.05), post-hoc contrasts were carried out between estimated means of model. Df, degrees of freedom; Df.res, Residual Degrees of Freedom; F, F-test statistic; p, p-value; x, not computed.

