## Supplementary Materials for "Alpha closed-loop auditory stimulation modulates waking alpha oscillations and sleep onset dynamics in a phase-dependent manner in humans"

### Supplementary Material

##### Table S1 | Phase-locking accuracy

Mean phase ($\phi_{mean}$) and standard deviation ($\phi_{SD}$) at stimulus onset, and target accuracy for each phase and location targeted in Study 1. The accuracy of phase locking was estimated using the Rayleigh test on the EEG data. Shown are the resultants (R), along with the z-stat (Z) and p values (p) from the Rayleigh test. **S1.1** – Data from the closed-loop EEG device. **S1.2** – Data from the high-density EEG device. Note that the two EEG systems are independent and different reference schemes were employed for the closed-loop and high-density EEG data.

| **S1.1 - Data from closed-loop EEG device** | | | | |
|---|---|---|---|---|
| | **Phase** | | | |
| **Location** | **Pre-Peak 330°** | **Post-Peak 60°** | **Pre-Trough 150°** | **Post-Trough 240°** |
| **Fz (N=20)** | $\phi_{mean}$ = 327.94°<br>$\phi_{SD}$ = 10.02°<br>$R_{mean}$ = 0.8466<br>$R_{SD}$ = 0.074<br>Z = 18.77<br>p = 1.27 × $10^{-13}$ | $\phi_{mean}$ = 53.31°<br>$\phi_{SD}$ = 10.04°<br>$R_{mean}$ = 0.8701<br>$R_{SD}$ = .078<br>Z = 18.83<br>p = 1.20 × $10^{-13}$ | $\phi_{mean}$ = 145.87°<br>$\phi_{SD}$ = 13.40°<br>$R_{mean}$ = 0.8286<br>$R_{SD}$ = 0.064<br>Z = 18.53<br>p = 5.21 × $10^{-13}$ | $\phi_{mean}$ = 237.69°<br>$\phi_{SD}$ = 12.60°<br>$R_{mean}$ = 0.8143<br>$R_{SD}$ = 0.078<br>Z = 18.51<br>p = 2.18 × $10^{-13}$ |
| **Pz (N=28)** | $\phi_{mean}$ = 333.32°<br>$\phi_{SD}$ = 9.82°<br>$R_{mean}$ = 0.8652<br>$R_{SD}$ = 0.047<br>Z = 18.77<br>p = 2.09 × $10^{-19}$ | $\phi_{mean}$ = 61.90°<br>$\phi_{SD}$ = 8.23°<br>$R_{mean}$ = 0.8432<br>$R_{SD}$ = 0.051<br>Z = 26.20<br>p = 2.55 × $10^{-19}$ | $\phi_{mean}$ = 155.122°<br>$\phi_{SD}$ = 10.95°<br>$R_{mean}$ = 0.8444<br>$R_{SD}$ = 0.053<br>Z = 25.99<br>p = 5.21 × $10^{-19}$ | $\phi_{mean}$ = 244.19°<br>$\phi_{SD}$ = 11.30°<br>$R_{mean}$ = 0.8491<br>$R_{SD}$ = 0.050<br>Z = 26.16<br>p = 3.69 × $10^{-19}$ |

| **S1.2 - Data from high-density EEG device** | | | | |
|---|---|---|---|---|
| | **Phase** | | | |
| **Location** | **Pre-Peak 330°** | **Post-Peak 60°** | **Pre-Trough 150°** | **Post-Trough 240°** |
| **Fz (N=20)** | $\phi_{mean}$ = 337.10°<br>$\phi_{SD}$ = 12.49°<br>$R_{mean}$ = 0.4897<br>$R_{SD}$ = 0.127<br>Z = 8.99<br>p = 1.53 × $10^{-5}$ | $\phi_{mean}$ = 72.04°<br>$\phi_{SD}$ = 10.96°<br>$R_{mean}$ = 0.5527<br>$R_{SD}$ = 0.134<br>Z = 10.07<br>p = 5.59 × $10^{--6}$ | $\phi_{mean}$ = 157.17°<br>$\phi_{SD}$ = 9.60°<br>$R_{mean}$ = 0.5757<br>$R_{SD}$ = 0.108<br>Z = 10.69<br>p = 2.45 × $10^{-6}$ | $\phi_{mean}$ = 240.07°<br>$\phi_{SD}$ = 10.02°<br>$R_{mean}$ = 0.5339<br>$R_{SD}$ = 0.134<br>Z = 9.45<br>p = 8.10 × $10^{-6}$ |
| **Pz (N=28)** | $\phi_{mean}$ = 345.32°<br>$\phi_{SD}$ = 15.27°<br>$R_{mean}$ = 0.6455 | $\phi_{mean}$ = 73.66°<br>$\phi_{SD}$ = 13.35<br>$R_{mean}$ = 0.6184 | $\phi_{mean}$ = 162.90°<br>$\phi_{SD}$ = 14.09°<br>$R_{mean}$ = 0.6456 | $\phi_{mean}$ = 251.43°<br>$\phi_{SD}$ = 13.98°<br>$R_{mean}$ = 0.6236 |

| | | | | |
|---|---|---|---|---|
| | $R_{SD} = 0.142$<br>$Z = 16.82$<br>$p = 1.12 \times 10^{-8}$ | $R_{SD} = 0.111$<br>$Z = 15.79$<br>$p = 2.33 \times 10^{-8}$ | $R_{SD} = 0.130$<br>$Z = 16.49$<br>$p = 1.17 \times 10^{-8}$ | $R_{SD} = 0.131$<br>$Z = 16.03$<br>$p = 1.67 \times 10^{-8}$ |

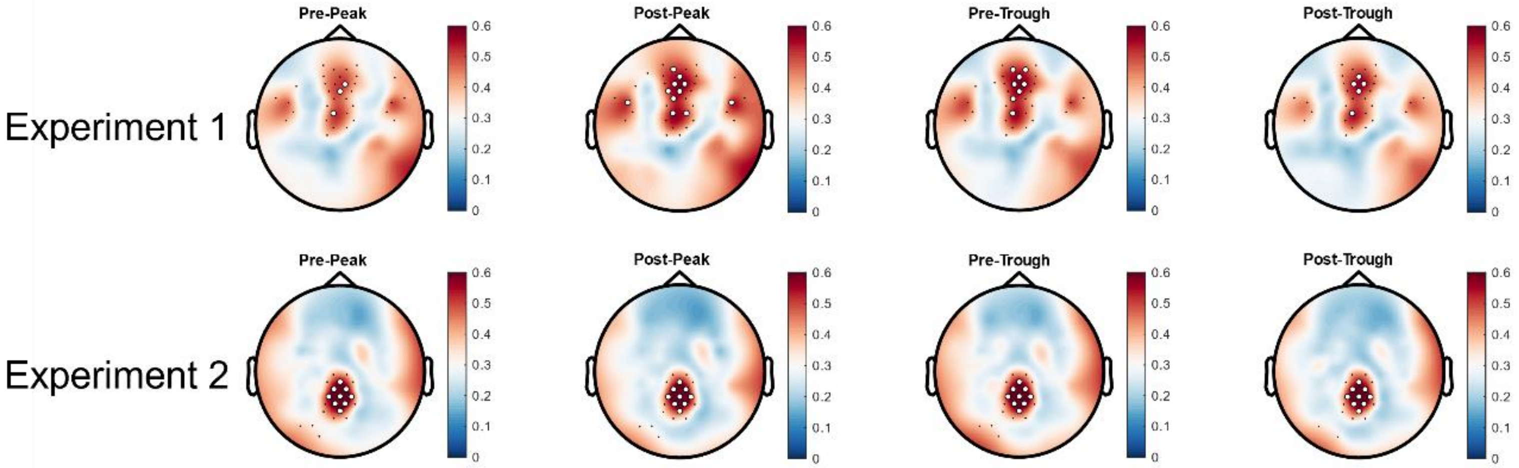


**Figure S1 Phase-locking Accuracy Across Conditions** – topography of phase-locking accuracy (average resultant) across 4 conditions of two experiments. White marks indicate channels at which resultant >0.5 and p <0.05. Black marks indicate channels at which resultant <0.5 and p <0.05. p values from Bonferroni-corrected z-test for non-uniformity.

**Figure S2 | power change ANOVA per 1 Hz band**

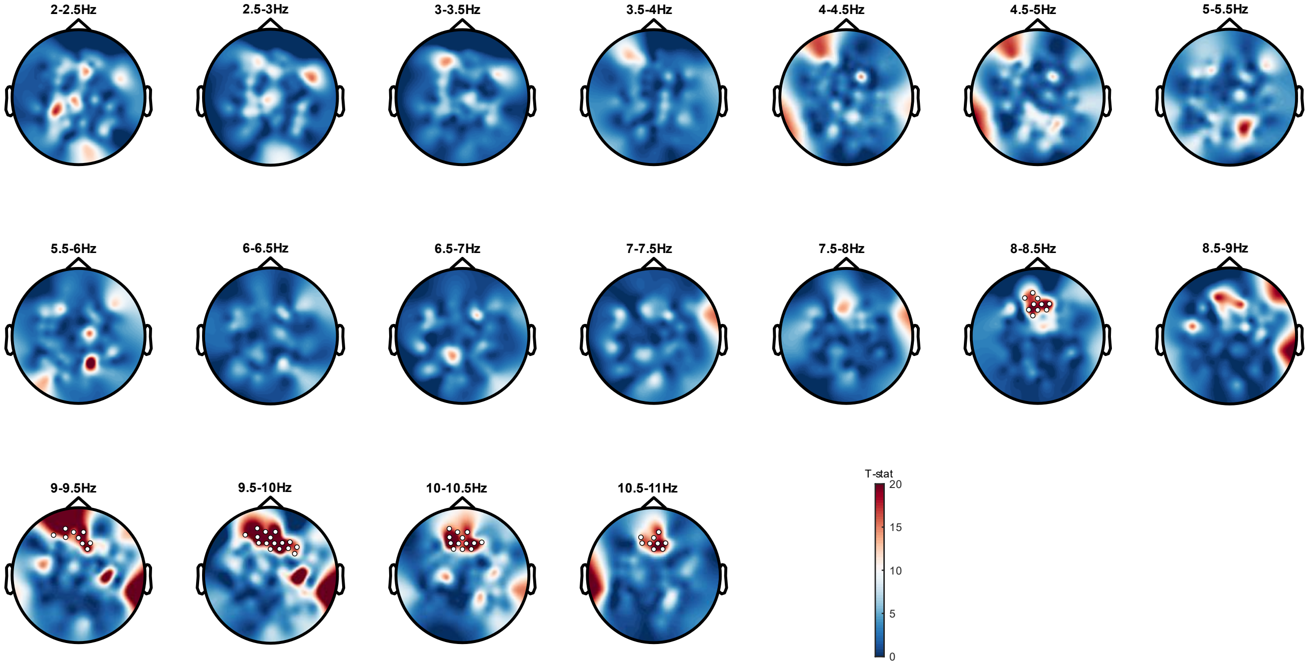


**Figure S2.1 Power change ANOVA –** topography of permutation ANOVA stats for each 0.5 Hz bin in experiment 1. White dots show significant main effect of phase-targeted, cluster-corrected *p* <0.05

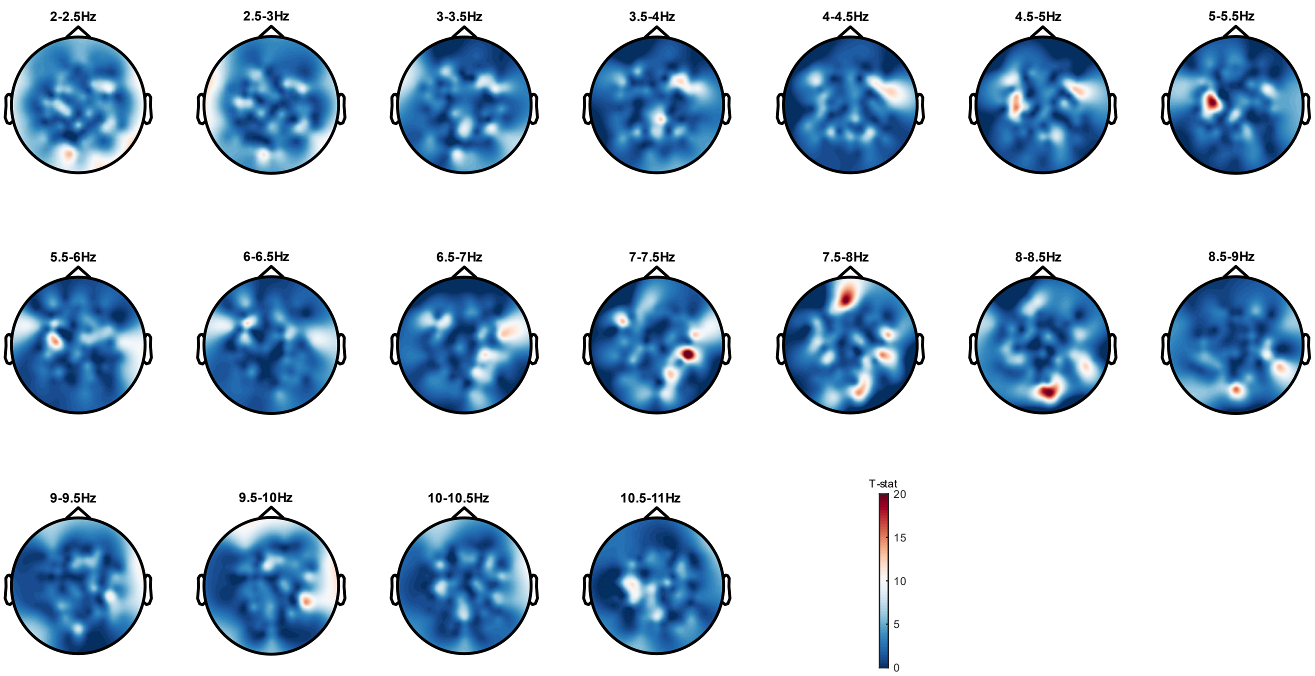


**Figure S2.2 Power change ANOVA** – topography of permutation ANOVA stats for each 0.5 Hz bin in experiment 1. White dots show significant main effect of phase-targeted, cluster-corrected *p* <0.05

**Figure S3 | power change within alpha band**

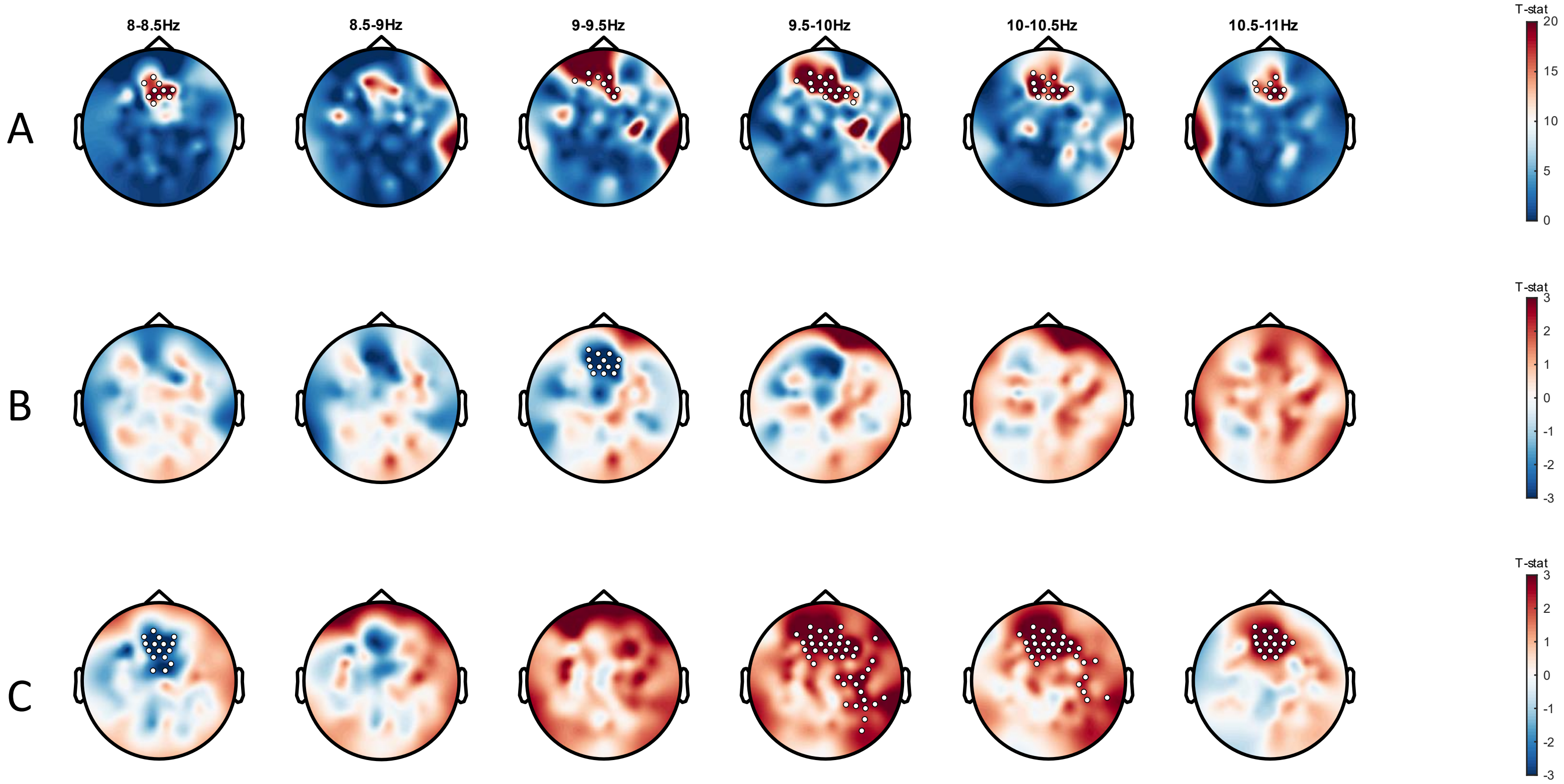


**Figure S3.1 Power change ANOVA** – (**A)** topography of permutation ANOVA stats for each .5 Hz bin in experiment 1. **(B)** pre-peak vs pre-trough, positive values indicate pre-trough is greater **(C)** post-peak vs post-trough, positive values indicate post-peak is greater. White dots show significant main effect of phase-targeted, cluster-corrected *p* <0.05.

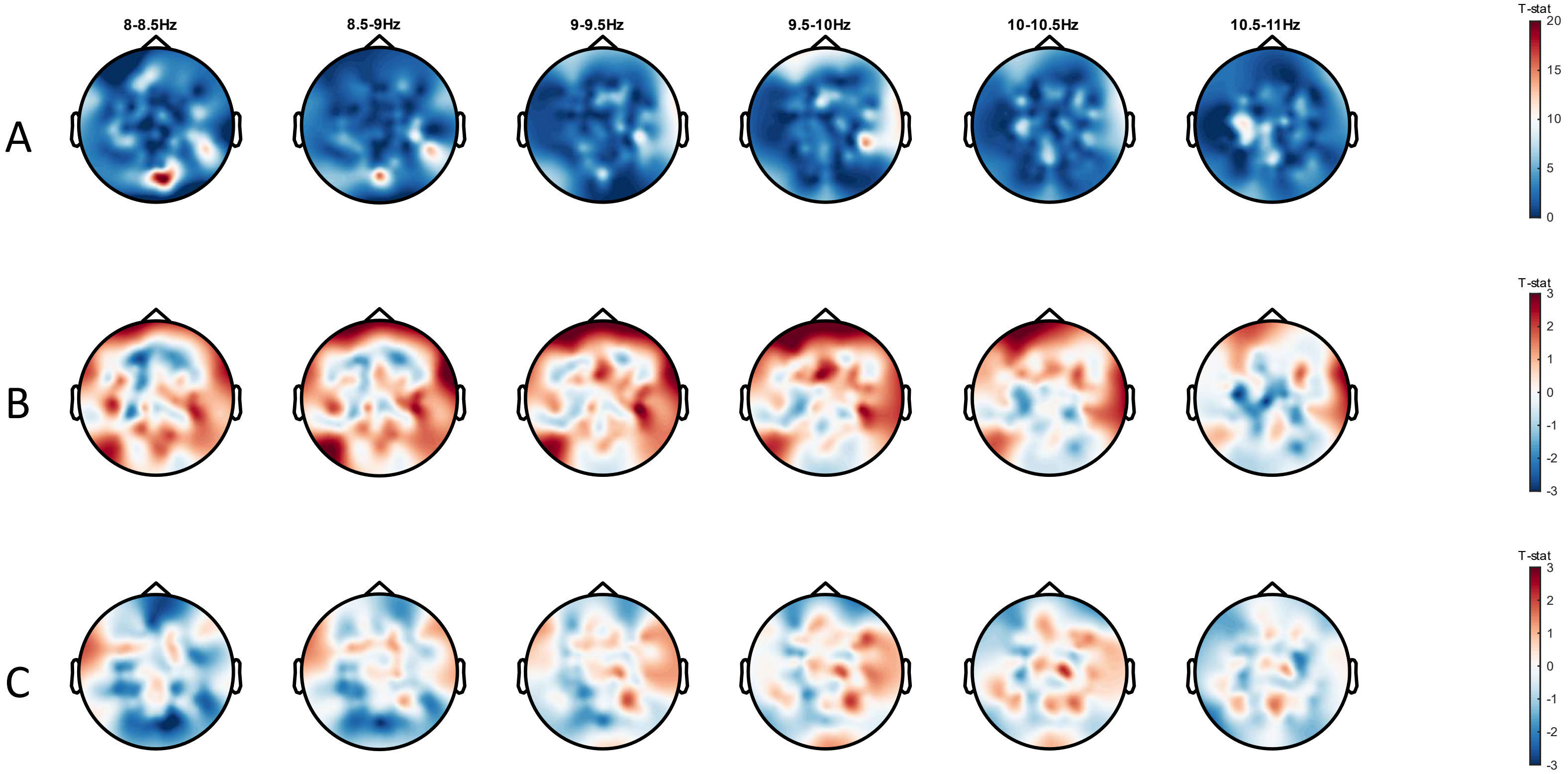


**Figure S3.2 Power change ANOVA** – (**A)** topography of permutation ANOVA stats for each 0.5 Hz bin in experiment 1. **(B)** pre-peak vs pre-trough, positive values indicate pre-trough is greater **(C)** post-peak vs post-trough, positive values indicate post-peak is greater. White dots show significant main effect of phase-targeted, cluster-corrected $p$ <0.05.

**Figure S4 | connectivity change per band**

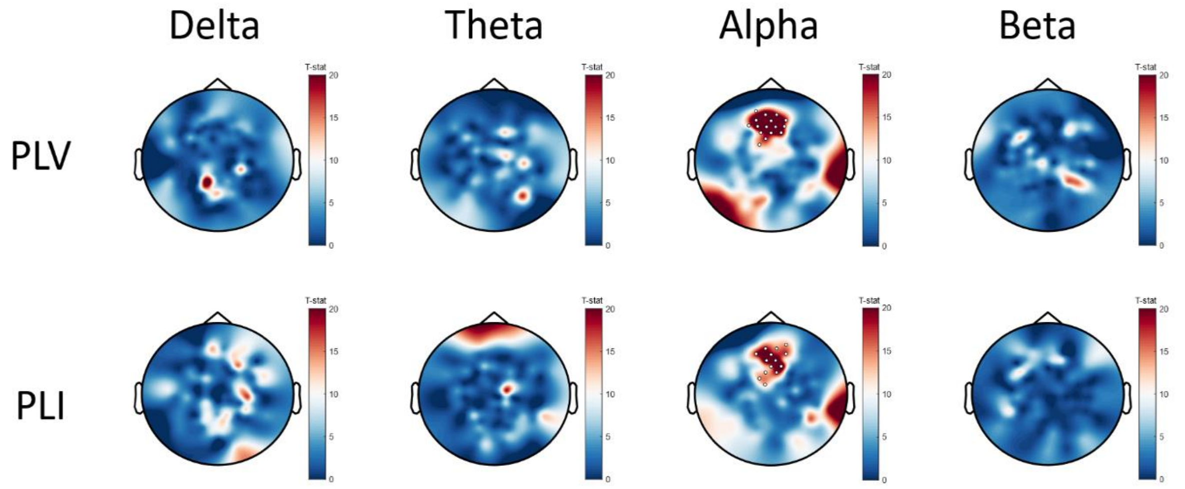


**Figure S4.1** topography of permutation ANOVA stats for each frequency band in experiment 1. White dots show significant main effect of phase-targeted, cluster-corrected $p < 0.05$.

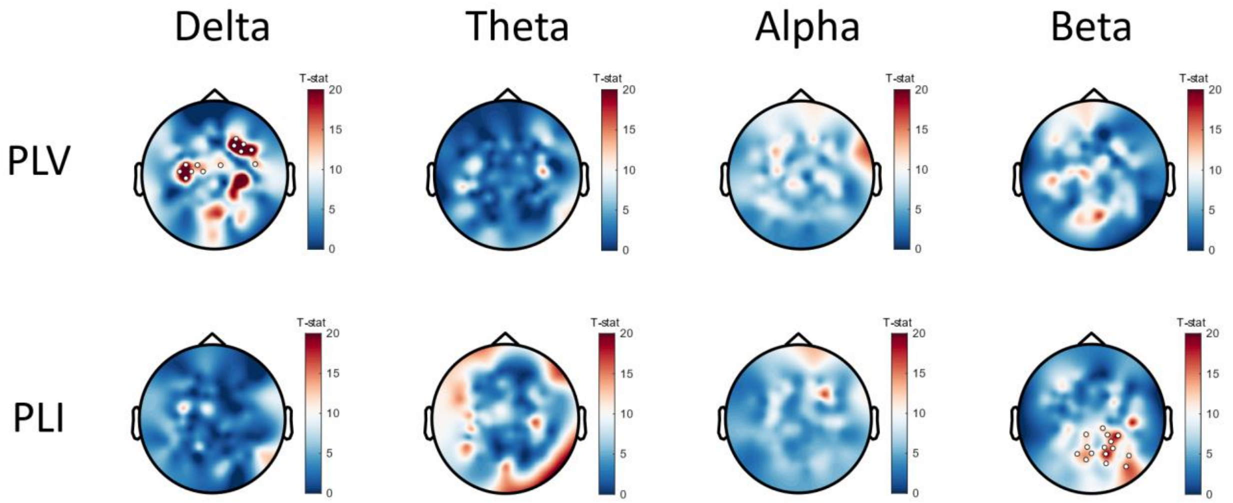


**Figure S4.2** topography of permutation ANOVA stats for each frequency band in experiment 2. White dots show significant main effect of phase-targeted, cluster-corrected *p* <0.05.

**Figure S5 | Alpha Frequency in hd-EEG and ecHT-EEG systems and Inter-Stimulus Intervals**

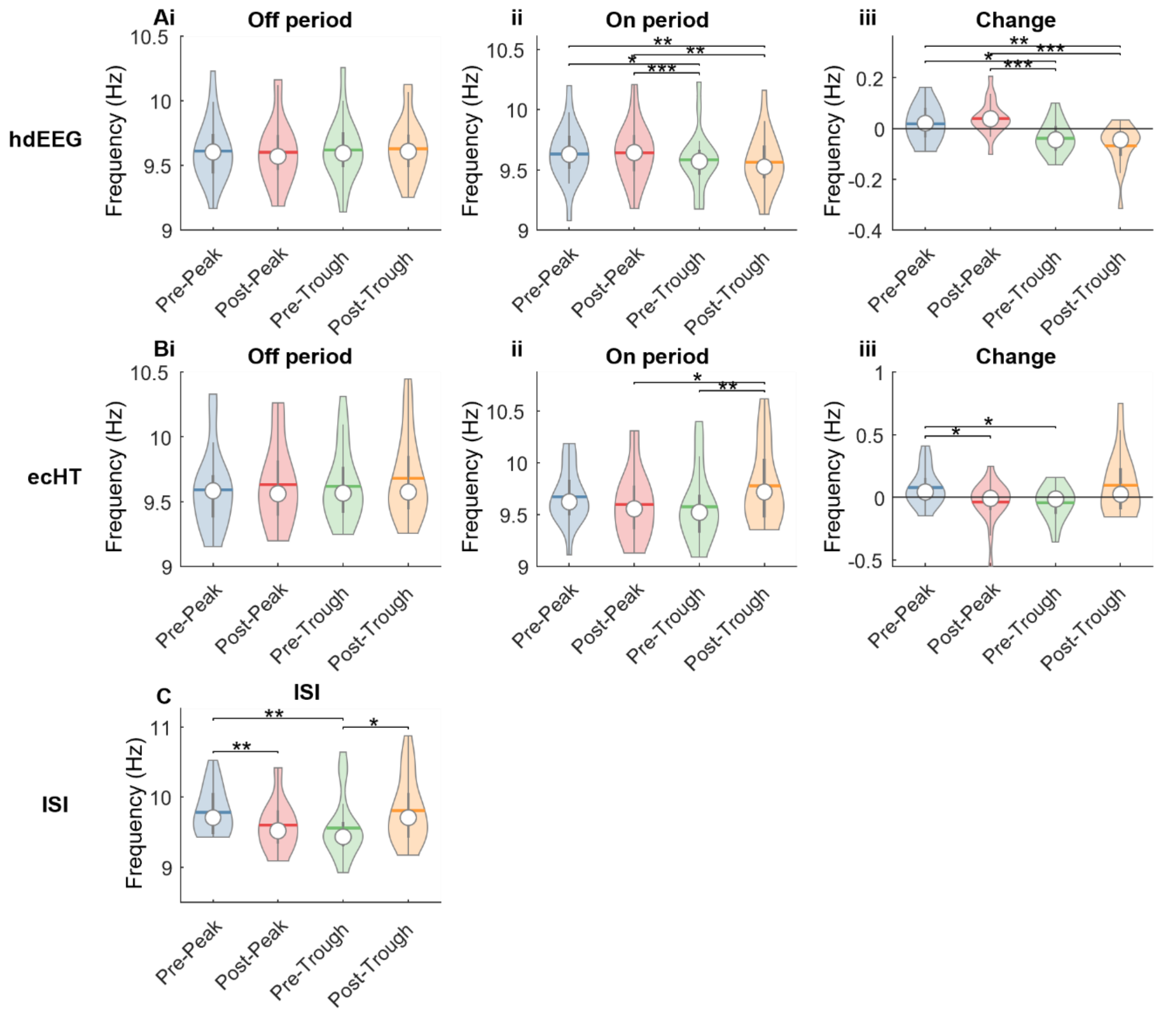


**Figure S5.1 (A)**Frequency estimates form frontal ROI in experiment 1; **(B)** Frequency estimates form ecHT in experiment 1 **(i)** Alpha frequency during 'Off' period; **(ii)** Alpha frequency during 'On' period; **(iii)** Sound-induced change in alpha frequency **(C)** Inter-Stimulus Interval. For all plots, mixed effects models were run: [frequency/ISI ~ condition + (1|Participant)]. Post-hoc t-tests were run for those which showed a statistically significant effect of condition ($p < .05$). * $p$<.05, ** $p$<.01, *** $p$<.001.

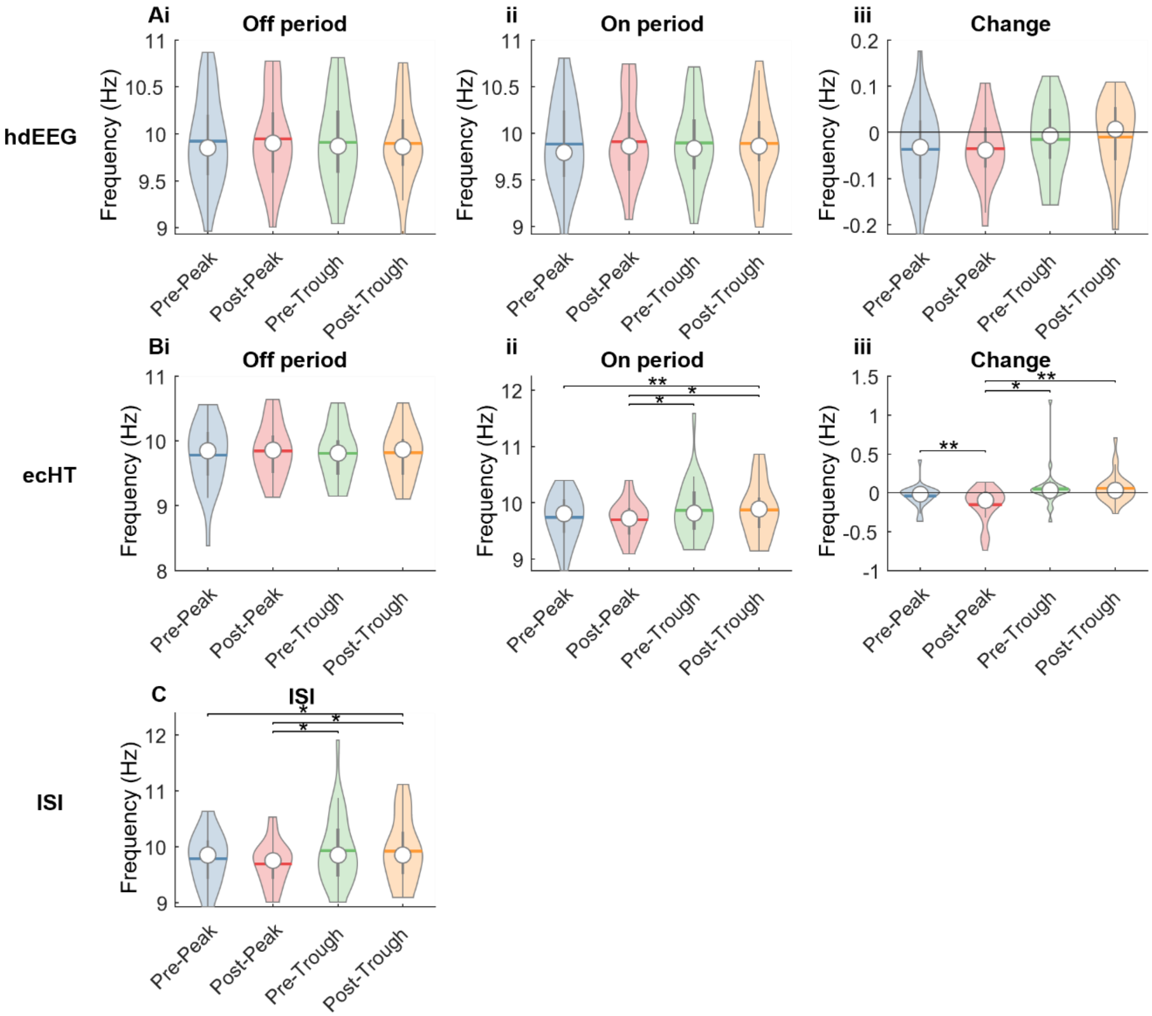


**Figure S5.2 (A)**Frequency estimates form parietal ROI in experiment 2; **(B)** Frequency estimates form ecHT in experiment 2 **(i)** Alpha frequency during 'Off' period; **(ii)** Alpha frequency during 'On' period; **(iii)** Sound-induced change in alpha frequency **(C)** Inter-Stimulus Interval. For all plots, mixed effects models were run: [frequency/ISI ~ condition + (1|Participant)]. Post-hoc t-tests were run for those which showed a statistically significant effect of condition ($p$ < .05). * $p$<.05, **$p$<.01, ***$p$<.001.

Whilst ISI did different between some conditions in both experiment 1 and 2, it did not seem to drive the frequency changes in the frontal ROI in experiment 1, as taken from the hdEEG, and instead appeared more consistent with the frequency estimates taken from the ecHT. Regardless, we go on to show that the stimulator frequency follows the brain activity, in study 3.

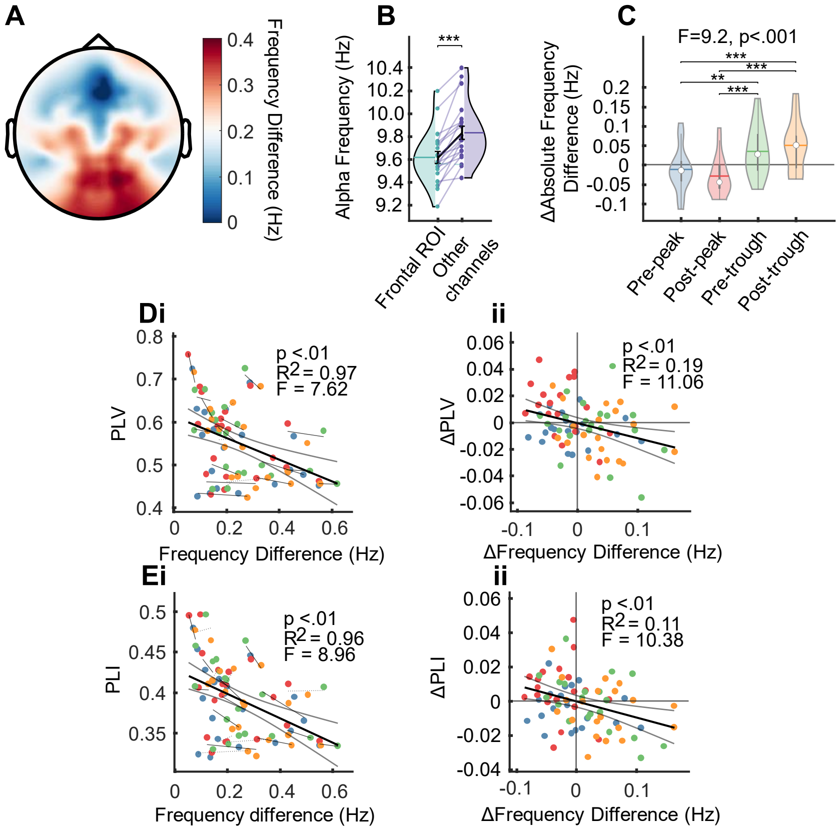


**Figure S6. Connectivity is partially accounted for by frequency difference in experiment 1**

**(A)** Topography of alpha frequency difference from frontal ROI during off (i.e. no stimulation) periods. **(B)** Alpha frequency at frontal ROI vs all other channels during off periods.† **(C)** Average stimulation-induced changes in alpha frequency differences, between frontal ROI and all other channels. †† **(D)** **(i)** Average frequency difference between frontal ROI and all other channels vs average PLV between frontal ROI and all other channels, lines indicate individual participant linear fit across four conditions††† **(ii)** Average stimulation-induced frequency difference change between frontal ROI and all other channels vs average stimulation-induced PLV change between frontal ROI and all other channels. Crossed lines indicate zero.††† **(E)(i)(ii)** same as **(D)** but for PLI†††.

†Stats are from a paired t-test, *** *p*<.001

††Stats are from a LMEM: ΔAbsolute_frequency_difference ~ Condition + (1|Participant)

††† Stats are from a LMEM; Connectivity ~ Frequency_difference + (1|Participant) **or** ΔConnectivity ~ ΔFrequency_difference + (1|Participant)

#### Figure S7 | Study 2 Resultant per Octile

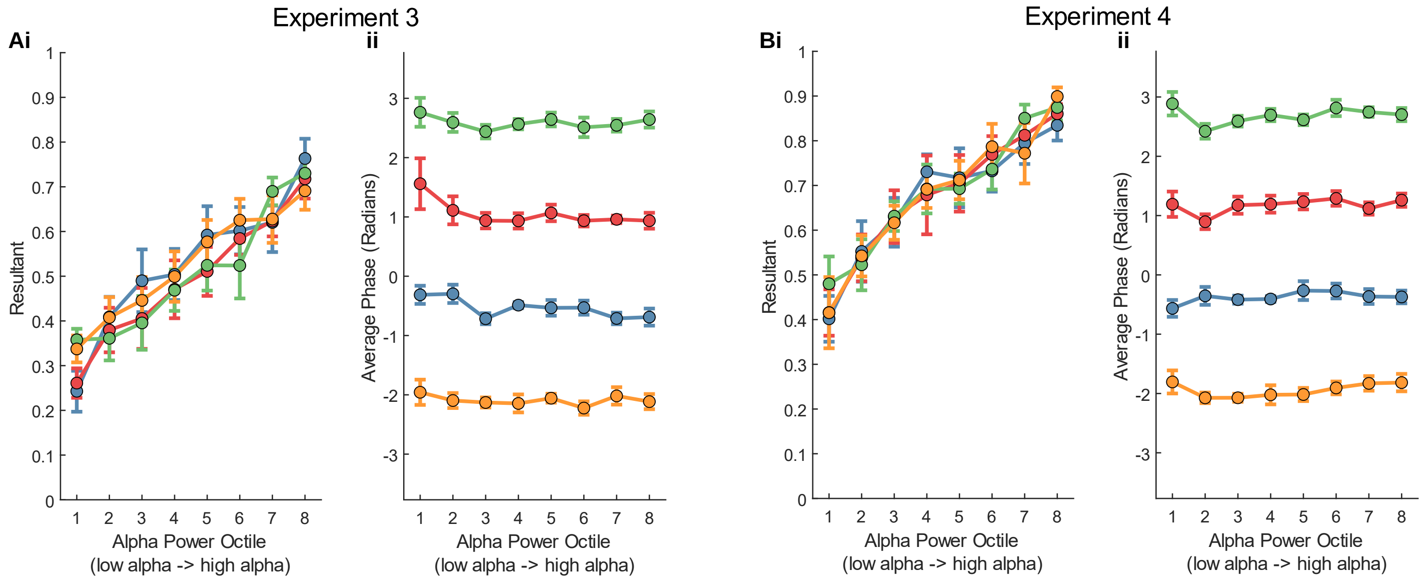


**Figure S7 Resultant per Octile. (A) (B) (C) (D)**

There was a clear linear relationship between alpha power octile and stimulus onset resultant in each condition, in both experiments (Figure SX). This meant that the phase was most consistent between octile 8 trials, and least consistent between octile 1 trials. We considered that this likely resulted from the use of two independent EEG systems, since the phase-locking (ecHT) system and the hdEEG will be in greatest agreement, regarding phase, when alpha power is high, and stimulus onset is determined only by the ecHT. We suggest that the extent of the reset should not be dependent on this onset resultant.

We tested this by taking 10,00 samples of 20 trials from each conditions, computing stimulus onset resultant, averaging across conditions, and plotting this against auditory-evoked Z statistic. We found a statistically significant, but very weak relationship, and hence confirmed our intuition that z-stat is not strongly dependent on onset resultant, and that alpha amplitude was

The average phase angle was highly consistent across octiles (Figure S8).

**Figure S8 | Study 2 Resultant vs Z Stat**

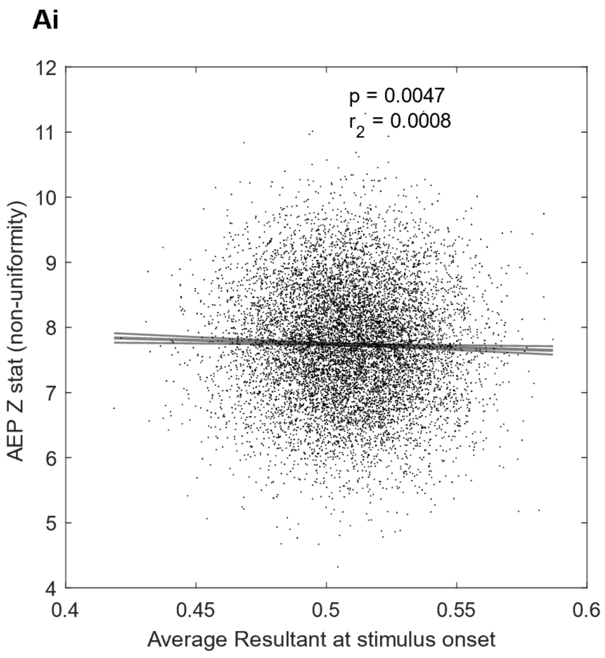


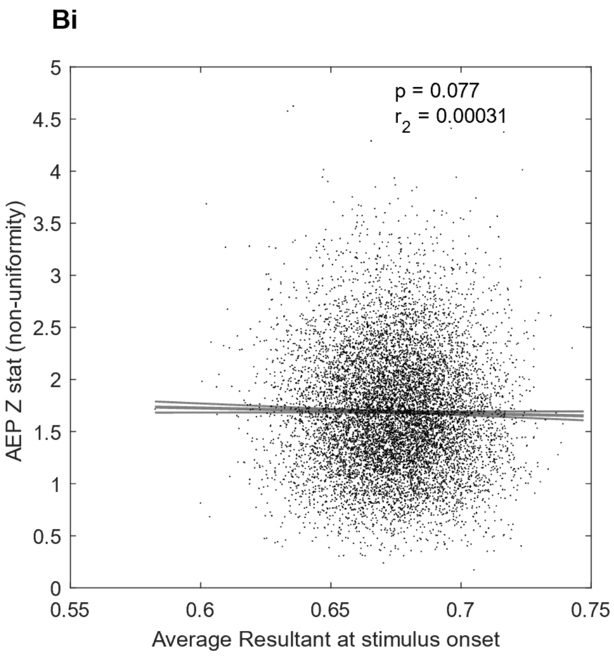

**S2 – Trials in each phase bin**

| Experiment/Location | | Phase (°) | | | | | | | | | |
|---|---|---|---|---|---|---|---|---|---|---|---|
| | | **0** | **36** | **72** | **108** | **144** | **180** | **216** | **252** | **288** | **324** |
| **3/Fz (N=8)** | **Mean** | 66.8 | 80 | 74.8 | 71.5 | 71.8 | 67.1 | 80 | 76.3 | 77.8 | 74.1 |
| | **SD** | 8.0 | 6.7 | 6.7 | 8.0 | 9.4 | 10.6 | 10.1 | 9.6 | 6.8 | 8.0 |
| **4/Pz (N=7)** | **Mean** | 76.9 | 67.1 | 80.4 | 67.4 | 74.3 | 70.1 | 71.7 | 69 | 68.1 | 76.9 |
| | **SD** | 14.1 | 17.4 | 8.4 | 11.8 | 12.7 | 17.8 | 14.2 | 12.0 | 14.0 | 14.1 |

**Figure S9 | Study 3 phase histograms**

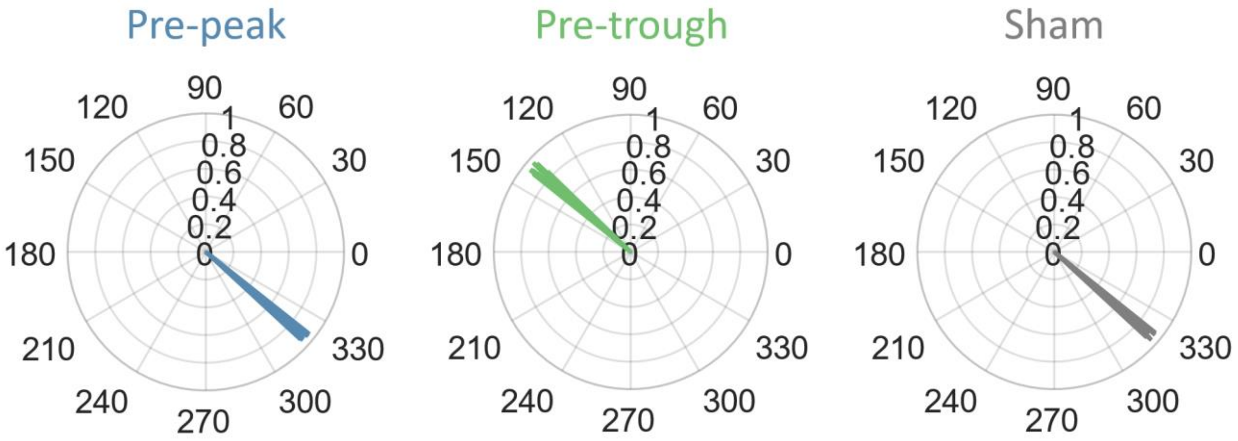


**Figure S9** Phase accuracy plots for the ecHT electrode in the three conditions in study 3. Each line represents a participant, length of line indicates resultant (between 0 and 1). Phase accuracy is high in all three conditions. Phases are the same for pre-peak and sham. During sham, markers were recorded for each sound stimulus but the volume was zero.

**Figure S10 – Study 3, percentage  of ISI’s per frequency band, per vigilance state**

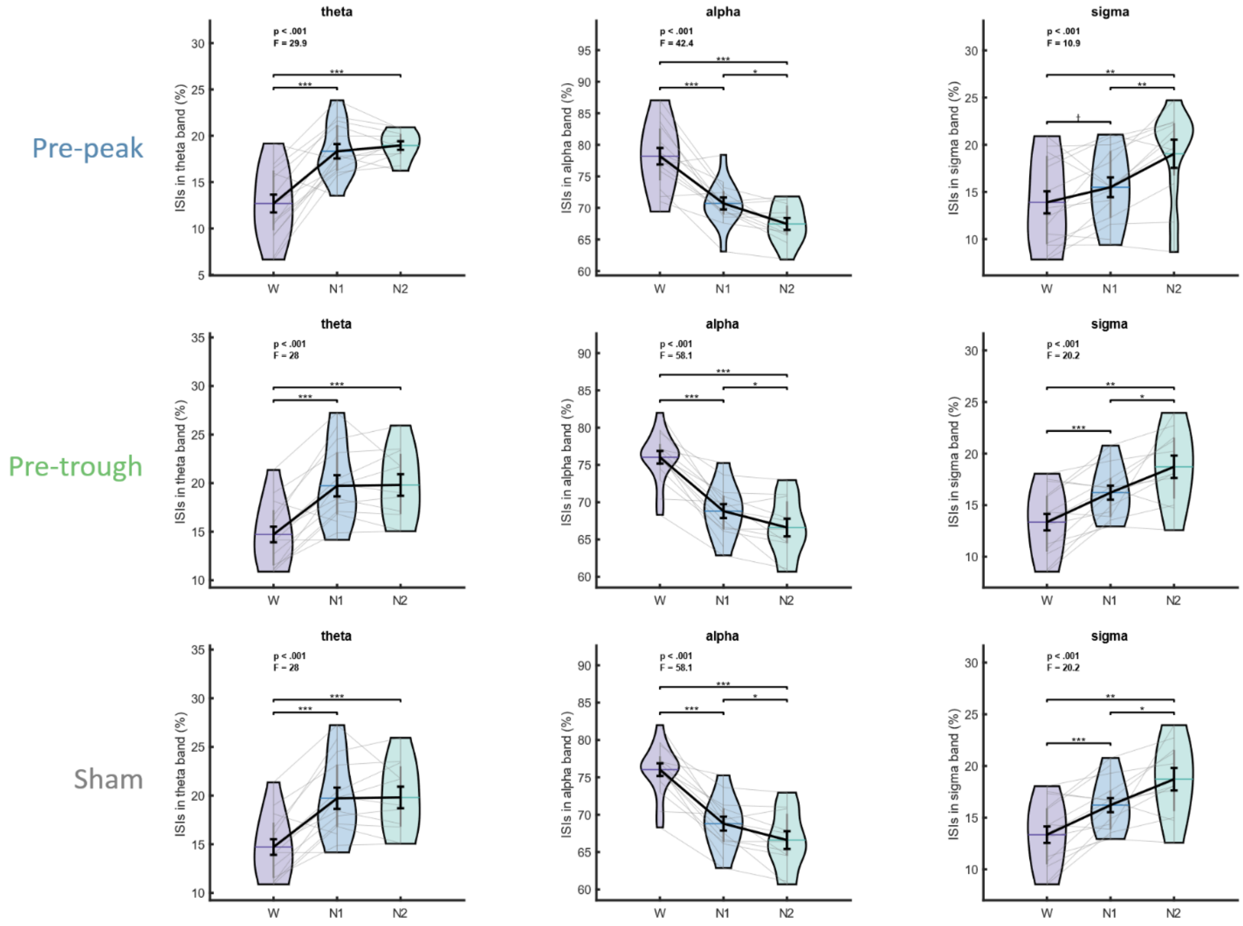


**Figure S10 – Inter stimulus intervals per sleep stage.** Violin plots show the percentage of ISI’s in each frequency band, in each sleep stage, in each condition. Stats indicate output of linear mixed effects model [ISI_percentage ~ sleep_stage + (1|participant)]. * p <.05, ** p<.01, ***p<.001.

#### Figure S11 – Average time series for all eBOSC features

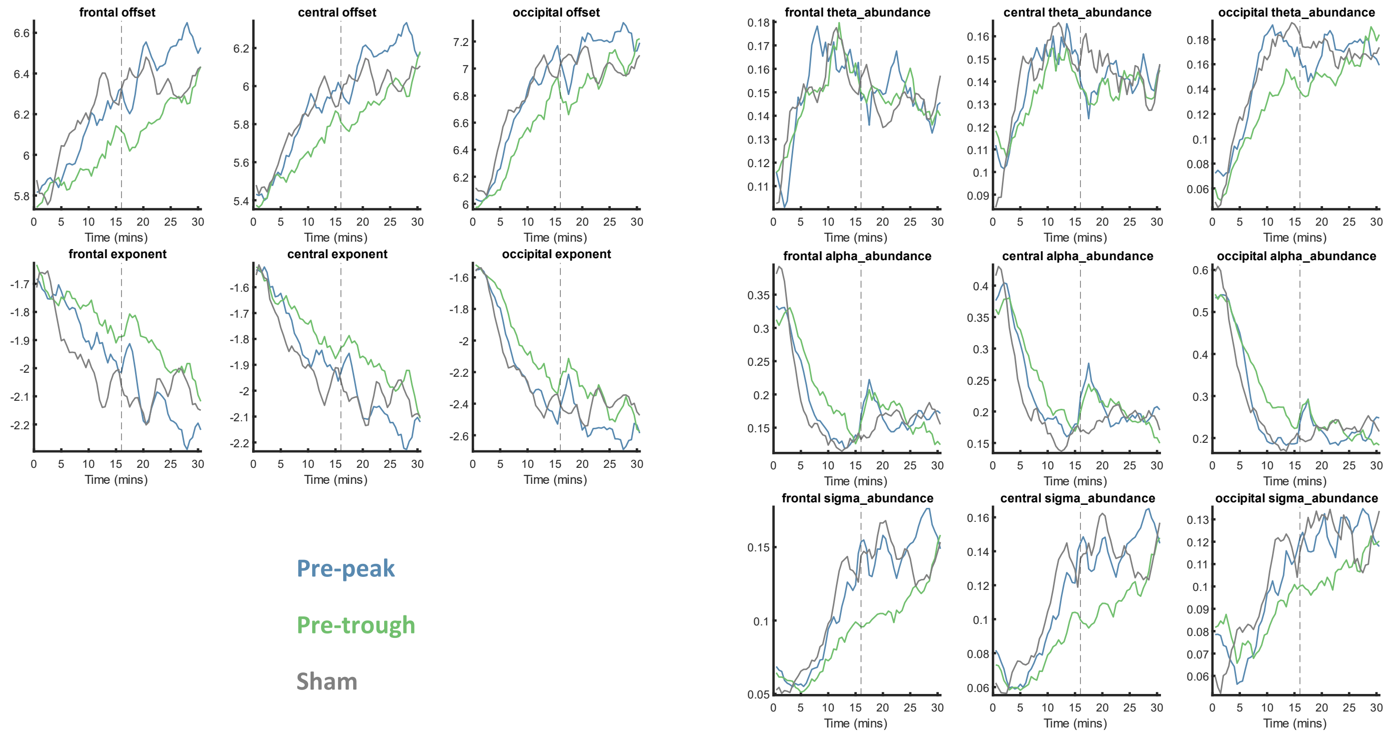


**Figure S11 – Time series for each eBOSC features.** Average time series are shown for each feature, each region, for each condition. Data was smoothed using a 2 minute moving mean window

#### Figure S12 – Average eBOSC features per sleep stage

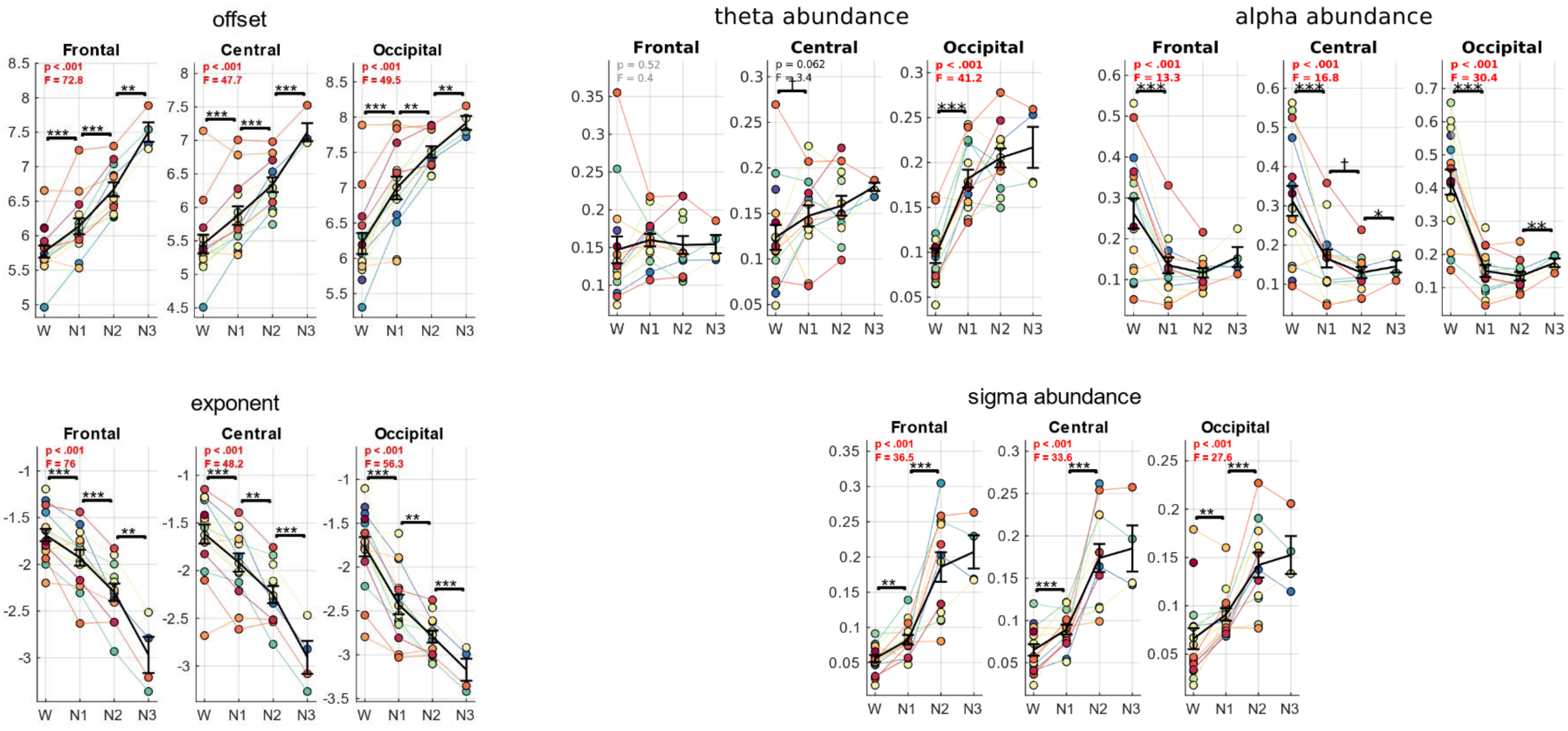


**Figure S12– eBOSC features per sleep stage.** Average are shown for each feature, each region, each participant, for the sham condition. Stats indicate output of linear mixed effects model [eBOSC_feature ~ sleep_stage + (1|participant)]. * p <.05, ** p<.01, ***p<.001.

#### Table S3 | Sleep scoring

Duration in minutes of each vigilance state for each participant, in each condition. Values are broken down into stimulation, post-stimulation, and whole nap periods.

| S2.1 sleep stage durations | | | | | | | | | | | | | |
|---|---|---|---|---|---|---|---|---|---|---|---|---|---|
| **Participant** | **Condition** | **Stim Wake** | **Stim N1** | **Stim N2** | **Stim N3** | **Post-Stim Wake** | **Post-Stim N1** | **Post-Stim N2** | **Post-Stim N3** | **Whole Nap Wake** | **Whole Nap N1** | **Whole Nap N2** | **Whole Nap N3** |
| **1** | Pre-Peak | 4.5 | 3 | 7.5 | 0 | 1 | 2.5 | 10.5 | 1 | 6.5 | 5.5 | 18 | 1 |
| | Post-Peak | 12 | 3 | 0 | 0 | 6 | 8 | 1 | 0 | 19 | 11 | 1 | 0 |
| | Sham | 15 | 0 | 0 | 0 | 15 | 0 | 0 | 0 | 31 | 0 | 0 | 0 |
| **2** | Pre-Peak | 3.5 | 3.5 | 8 | 0 | 0.5 | 1.5 | 9 | 4 | 5 | 5 | 17 | 4 |
| | Post-Peak | 5.5 | 3 | 6.5 | 0 | 1 | 5 | 9 | 0 | 7.5 | 8 | 15.5 | 0 |
| | Sham | 3.5 | 2.5 | 9 | 0 | 6 | 3.5 | 5 | 0.5 | 10.5 | 6 | 14 | 0.5 |
| **3** | Pre-Peak | 6.5 | 2 | 6.5 | 0 | 3.5 | 1.5 | 10 | 0 | 11 | 3.5 | 16.5 | 0 |
| | Post-Peak | 13.5 | 1.5 | 0 | 0 | 6 | 8.5 | 0.5 | 0 | 20.5 | 10 | 0.5 | 0 |
| | Sham | 5 | 4 | 6 | 0 | 0 | 0.5 | 14.5 | 0 | 6 | 4.5 | 20.5 | 0 |
| **4** | Pre-Peak | 2 | 1 | 4 | 8 | 0 | 0 | 0.5 | 14.5 | 3 | 1 | 4.5 | 22.5 |
| | Post-Peak | 4.5 | 8.5 | 2 | 0 | 0 | 0.5 | 14.5 | 0 | 5.5 | 9 | 16.5 | 0 |
| | Sham | 3 | 4 | 8 | 0 | 0 | 0 | 8 | 7 | 4 | 4 | 16 | 7 |
| **5** | Pre-Peak | 4 | 4 | 7 | 0 | 3 | 4 | 8 | 0 | 8 | 8 | 15 | 0 |
| | Post-Peak | 12 | 2 | 1 | 0 | 3 | 6 | 6 | 0 | 16 | 8 | 7 | 0 |
| | Sham | 4.5 | 4.5 | 6 | 0 | 1 | 5 | 9 | 0 | 6.5 | 9.5 | 15 | 0 |
| **6** | Pre-Peak | 15 | 0 | 0 | 0 | 6.5 | 8.5 | 0 | 0 | 22.5 | 8.5 | 0 | 0 |
| | Post-Peak | 8 | 7 | 0 | 0 | 1.5 | 9.5 | 4 | 0 | 10.5 | 16.5 | 4 | 0 |
| | Sham | 8.5 | 4.5 | 2 | 0 | 1 | 3.5 | 10.5 | 0 | 10.5 | 8 | 12.5 | 0 |
| **7** | Pre-Peak | 13 | 2 | 0 | 0 | 4.5 | 1.5 | 9 | 0 | 18.5 | 3.5 | 9 | 0 |
| | Post-Peak | 13 | 2 | 0 | 0 | 2.5 | 5 | 7.5 | 0 | 16.5 | 7 | 7.5 | 0 |
| | Sham | 10 | 5 | 0 | 0 | 0 | 1.5 | 13.5 | 0 | 11 | 6.5 | 13.5 | 0 |
| **8** | Pre-Peak | 11.5 | 3.5 | 0 | 0 | 5 | 6.5 | 3.5 | 0 | 17.5 | 10 | 3.5 | 0 |
| | Post-Peak | 13 | 2 | 0 | 0 | 5.5 | 4.5 | 5 | 0 | 19.5 | 6.5 | 5 | 0 |
| | Sham | 13.5 | 1.5 | 0 | 0 | 11 | 3.5 | 0.5 | 0 | 25.5 | 5 | 0.5 | 0 |
| **9** | Pre-Peak | 6 | 5.5 | 3.5 | 0 | 0 | 0.5 | 7 | 7.5 | 7 | 6 | 10.5 | 7.5 |
| | Post-Peak | 12.5 | 1.5 | 1 | 0 | 1.5 | 7 | 6.5 | 0 | 15 | 8.5 | 7.5 | 0 |
| | Sham | 6 | 2.5 | 6.5 | 0 | 0.5 | 2.5 | 9.5 | 2.5 | 7.5 | 5 | 16 | 2.5 |
| **10** | Pre-Peak | 7.5 | 4 | 3.5 | 0 | 8.5 | 0 | 6.5 | 0 | 17 | 4 | 10 | 0 |
| | Post-Peak | 2.5 | 5.5 | 7 | 0 | 3 | 5.5 | 6.5 | 0 | 6.5 | 11 | 13.5 | 0 |
| | Sham | 4 | 6.5 | 4.5 | 0 | 3.5 | 2.5 | 9 | 0 | 8.5 | 9 | 13.5 | 0 |
| **11** | Pre-Peak | 15 | 0 | 0 | 0 | 9.5 | 5 | 0.5 | 0 | 25.5 | 5 | 0.5 | 0 |
| | Post-Peak | 11.5 | 3.5 | 0 | 0 | 13.5 | 1.5 | 0 | 0 | 26 | 5 | 0 | 0 |
| | Sham | 14.5 | 0.5 | 0 | 0 | 14.5 | 0.5 | 0 | 0 | 30 | 1 | 0 | 0 |
| **12** | Pre-Peak | 4.5 | 2.5 | 8 | 0 | 0.5 | 3.5 | 11 | 0 | 6 | 6 | 19 | 0 |
| | Post-Peak | 6 | 3 | 6 | 0 | 4.5 | 3 | 7.5 | 0 | 11.5 | 6 | 13.5 | 0 |
| | Sham | 4.5 | 6 | 4.5 | 0 | 6 | 4.5 | 4.5 | 0 | 11.5 | 10.5 | 9 | 0 |
| **13** | Pre-Peak | 3.5 | 2.5 | 9 | 0 | 0 | 0 | 4 | 11 | 4.5 | 2.5 | 13 | 11 |
| | Post-Peak | 5.5 | 4 | 5.5 | 0 | 0 | 0 | 2.5 | 12.5 | 6.5 | 4 | 8 | 12.5 |
| | Sham | 2 | 3.5 | 9 | 0.5 | 0 | 0 | 1.5 | 13.5 | 3 | 3.5 | 10.5 | 14 |
| **14** | Pre-Peak | 4.5 | 4 | 6.5 | 0 | 4.5 | 4.5 | 6 | 0 | 10 | 8.5 | 12.5 | 0 |
| | Post-Peak | 4.5 | 10.5 | 0 | 0 | 9 | 3.5 | 2.5 | 0 | 14.5 | 14 | 2.5 | 0 |
| | Sham | 5 | 4.5 | 5.5 | 0 | 0 | 2 | 13 | 0 | 6 | 6.5 | 18.5 | 0 |
| **15** | Pre-Peak | 5.5 | 9 | 0.5 | 0 | 3.5 | 3.5 | 4.5 | 3.5 | 10 | 12.5 | 5 | 3.5 |
| | Post-Peak | 4.5 | 10 | 0.5 | 0 | 4 | 8 | 3 | 0 | 9.5 | 18 | 3.5 | 0 |
| | Sham | 6.5 | 4.5 | 4 | 0 | 8.5 | 4 | 2.5 | 0 | 16 | 8.5 | 6.5 | 0 |
| **16** | Pre-Peak | 14 | 1 | 0 | 0 | 14.5 | 0.5 | 0 | 0 | 29.5 | 1.5 | 0 | 0 |
| | Post-Peak | 15 | 0 | 0 | 0 | 10 | 5 | 0 | 0 | 26 | 5 | 0 | 0 |
| | Sham | 15 | 0 | 0 | 0 | 15 | 0 | 0 | 0 | 31 | 0 | 0 | 0 |

**Table S4 | eBOSC statistics**

Tables show results of linear mixed-effects models of the form:

[eBOSC_feature ~ condition + (1|participant)]

Where a main effect of condition was found (*p* <.05), post-hoc contrasts were carried out between estimated means of model

| S4.1 eBOSC statistics from stimulation period | | | | | |
|---|---|---|---|---|---|
| | Df | Df.res | F | p | |
| 'frontal_offset' | 2 | 45 | 5.923954 | 0.005202 | |
| Post-hoc contrasts | B | SE | df | t | p |
| pre_peak - pre_trough | 0.127973 | 6.063338 | 45 | 2.206821 | 0.03247 |
| pre_peak - sham | -0.06867 | 6.063338 | 45 | 1.184249 | 0.24253 |
| pre_trough - sham | -0.19665 | 6.063338 | 45 | 3.39107 | 0.00146 |
| | Df | Df.res | F | p | |
| 'frontal_exponent' | 2 | 45 | 6.924841 | 0.002388 | |
| Post-hoc contrasts | B | SE | df | t | p |
| pre_peak - pre_trough | -0.06163 | -1.85594 | 45 | 1.530077 | 0.133 |
| pre_peak - sham | 0.087522 | -1.85594 | 45 | 2.172888 | 0.03509 |
| pre_trough - sham | 0.149152 | -1.85594 | 45 | 3.702964 | 0.00058 |
| | Df | Df.res | F | p | |
| 'frontal_theta_abundance' | 2 | 45 | 0.156121 | 0.855917 | |
| Post-hoc contrasts | B | SE | df | t | p |
| pre_peak - pre_trough | X | X | X | X | X |
| pre_peak - sham | X | X | X | X | X |
| pre_trough - sham | X | X | X | X | X |
| | Df | Df.res | F | p | |
| 'frontal_alpha_abundance' | 2 | 45 | 5.471691 | 0.007463 | |
| Post-hoc contrasts | B | SE | df | t | p |
| pre_peak - pre_trough | -0.03095 | 0.184084 | 45 | 2.338787 | 0.02385 |
| pre_peak - sham | 0.011338 | 0.184084 | 45 | 0.856709 | 0.39615 |
| pre_trough - sham | 0.042291 | 0.184084 | 45 | 3.195496 | 0.00255 |
| | Df | Df.res | F | p | |
| 'frontal_sigma_abundance' | 2 | 45 | 6.631643 | 0.002992 | |
| Post-hoc contrasts | B | SE | df | t | p |
| pre_peak - pre_trough | 0.016751 | 0.088352 | 45 | 2.58685 | 0.01299 |
| pre_peak - sham | -0.006 | 0.088352 | 45 | 0.926625 | 0.35906 |
| pre_trough - sham | -0.02275 | 0.088352 | 45 | 3.513475 | 0.00102 |
| | Df | Df.res | F | p | |
| 'central_offset' | 2 | 45 | 3.873805 | 0.028036 | |
| Post-hoc contrasts | B | SE | df | t | p |
| pre_peak - pre_trough | 0.131816 | 5.75134 | 45 | 2.146797 | 0.03724 |
| pre_peak - sham | -0.0283 | 5.75134 | 45 | 0.460928 | 0.64707 |
| pre_trough - sham | -0.16012 | 5.75134 | 45 | 2.607724 | 0.01232 |
| | Df | Df.res | F | p | |
| 'central_exponent' | 2 | 45 | 4.061698 | 0.023897 | |
| Post-hoc contrasts | B | SE | df | t | p |
| pre_peak - pre_trough | -0.05631 | -1.78099 | 45 | 1.270906 | 0.21029 |
| pre_peak - sham | 0.069729 | -1.78099 | 45 | 1.573878 | 0.12252 |
| pre_trough - sham | 0.126035 | -1.78099 | 45 | 2.844784 | 0.00667 |
| | Df | Df.res | F | p | |
| 'central_theta_abundance' | 2 | 45 | 1.547957 | 0.223794 | |
| Post-hoc contrasts | B | SE | df | t | p |
| pre_peak - pre_trough | X | X | X | X | X |
| pre_peak - sham | X | X | X | X | X |
| pre_trough - sham | X | X | X | X | X |
| | Df | Df.res | F | p | |
| 'central_alpha_abundance' | 2 | 45 | 3.156508 | 0.052139 | |
| Post-hoc contrasts | B | SE | df | t | p |
| pre_peak - pre_trough | X | X | X | X | X |
| pre_peak - sham | X | X | X | X | X |
| pre_trough - sham | X | X | X | X | X |
| | Df | Df.res | F | p | |

| | | | | | |
|---|---|---|---|---|---|
| 'central_sigma_abundance' | 2 | 45 | 8.475196 | 0.000752 | |
| Post-hoc contrasts | B | SE | df | t | p |
| pre_peak - pre_trough | 0.015222 | 0.091094 | 45 | 2.732606 | 0.00895 |
| pre_peak - sham | -0.00724 | 0.091094 | 45 | 1.300613 | 0.20001 |
| pre_trough - sham | -0.02247 | 0.091094 | 45 | 4.033219 | 0.00021 |
| | Df | Df.res | F | p | |
| occipital_offset' | 2 | 45 | 3.275011 | 0.047004 | |
| Post-hoc contrasts | B | SE | df | t | p |
| pre_peak - pre_trough | 0.206954 | 6.649754 | 45 | 1.977196 | 0.05417 |
| pre_peak - sham | -0.04383 | 6.649754 | 45 | 0.418718 | 0.67741 |
| pre_trough - sham | -0.25078 | 6.649754 | 45 | 2.395914 | 0.0208 |
| | Df | Df.res | F | p | |
| occipital_exponent' | 2 | 45 | 3.068049 | 0.056352 | |
| Post-hoc contrasts | B | SE | df | t | p |
| pre_peak - pre_trough | X | X | X | X | X |
| pre_peak - sham | X | X | X | X | X |
| pre_trough - sham | X | X | X | X | X |
| | Df | Df.res | F | p | |
| occipital_theta_abundance' | 2 | 45 | 10.55111 | 0.000175 | |
| Post-hoc contrasts | B | SE | df | t | p |
| pre_peak - pre_trough | 0.030612 | 0.144317 | 45 | 3.892157 | 0.00033 |
| pre_peak - sham | -0.00131 | 0.144317 | 45 | 0.166978 | 0.86814 |
| pre_trough - sham | -0.03193 | 0.144317 | 45 | 4.059135 | 0.00019 |
| | Df | Df.res | F | p | |
| 'occipital_alpha_abundance' | 2 | 45 | 2.793389 | 0.071854 | |
| Post-hoc contrasts | B | SE | df | t | p |
| pre_peak - pre_trough | X | X | X | X | X |
| pre_peak - sham | X | X | X | X | X |
| pre_trough - sham | X | X | X | X | X |
| | Df | Df.res | F | p | |
| 'occipital_sigma_abundance' | 2 | 45 | 0.888401 | 0.418403 | |
| Post-hoc contrasts | B | SE | df | t | p |
| pre_peak - pre_trough | X | X | X | X | X |
| pre_peak - sham | X | X | X | X | X |
| pre_trough - sham | X | X | X | X | X |

#### S4.2 eBOSC statistics from post-stimulation period

| | | | | | |
|---|---|---|---|---|---|
| | Df | Df.res | F | p | |
| 'frontal_offset' | 2 | 45 | 3.252743 | 0.047927 | |
| Post-hoc contrasts | B | SE | df | t | p |
| pre_peak - pre_trough | 0.265652 | 6.473472 | 45 | 2.534944 | 0.01479 |
| pre_peak - sham | 0.107229 | 6.473472 | 45 | 1.02322 | 0.31167 |
| pre_trough - sham | -0.15842 | 6.473472 | 45 | 1.511724 | 0.1376 |
| | Df | Df.res | F | p | |
| 'frontal_exponent' | 2 | 45 | 3.12432 | 0.053633 | |
| Post-hoc contrasts | B | SE | df | t | p |
| pre_peak - pre_trough | X | X | X | X | X |
| pre_peak - sham | X | X | X | X | X |
| pre_trough - sham | X | X | X | X | X |
| | Df | Df.res | F | p | |
| 'frontal_theta_abundance' | 2 | 45 | 0.013075 | 0.987014 | |
| Post-hoc contrasts | B | SE | df | t | p |
| pre_peak - pre_trough | X | X | X | X | X |
| pre_peak - sham | X | X | X | X | X |
| pre_trough - sham | X | X | X | X | X |
| | Df | Df.res | F | p | |
| 'frontal_alpha_abundance' | 2 | 45 | 0.076834 | 0.926165 | |
| Post-hoc contrasts | B | SE | df | t | p |
| pre_peak - pre_trough | X | X | X | X | X |
| pre_peak - sham | X | X | X | X | X |
| pre_trough - sham | X | X | X | X | X |
| | Df | Df.res | F | p | |
| 'frontal_sigma_abundance' | 2 | 45 | 2.248611 | 0.117277 | |
| Post-hoc contrasts | B | SE | df | t | p |
| pre_peak - pre_trough | X | X | X | X | X |
| pre_peak - sham | X | X | X | X | X |
| pre_trough - sham | X | X | X | X | X |
| | Df | Df.res | F | p | |
| 'central_offset' | 2 | 45 | 2.316259 | 0.110291 | |
| Post-hoc contrasts | B | SE | df | t | p |
| pre_peak - pre_trough | X | X | X | X | X |
| pre_peak - sham | X | X | X | X | X |
| pre_trough - sham | X | X | X | X | X |
| | Df | Df.res | F | p | |
| 'central_exponent' | 2 | 45 | 1.913945 | 0.159313 | |
| Post-hoc contrasts | B | SE | df | t | p |
| pre_peak - pre_trough | X | X | X | X | X |
| pre_peak - sham | X | X | X | X | X |

| | | | | | |
|---|---|---|---|---|---|
| pre_trough - sham | X | X | X | X | X |
| | Df | Df.res | F | p | |
| 'central_theta_abundance' | 2 | 45 | 0.730922 | 0.487093 | |
| Post-hoc contrasts | B | SE | df | t | p |
| pre_peak - pre_trough | X | X | X | X | X |
| pre_peak - sham | X | X | X | X | X |
| pre_trough - sham | X | X | X | X | X |
| | Df | Df.res | F | p | |
| 'central_alpha_abundance' | 2 | 45 | 0.442127 | 0.64543 | |
| Post-hoc contrasts | B | SE | df | t | p |
| pre_peak - pre_trough | X | X | X | X | X |
| pre_peak - sham | X | X | X | X | X |
| pre_trough - sham | X | X | X | X | X |
| | Df | Df.res | F | p | |
| 'central_sigma_abundance' | 2 | 45 | 2.266612 | 0.115374 | |
| Post-hoc contrasts | B | SE | df | t | p |
| pre_peak - pre_trough | X | X | X | X | X |
| pre_peak - sham | X | X | X | X | X |
| pre_trough - sham | X | X | X | X | X |
| | Df | Df.res | F | p | |
| occipital_offset' | 2 | 45 | 1.245716 | 0.297467 | |
| Post-hoc contrasts | B | SE | df | t | p |
| pre_peak - pre_trough | X | X | X | X | X |
| pre_peak - sham | X | X | X | X | X |
| pre_trough - sham | X | X | X | X | X |
| | Df | Df.res | F | p | |
| occipital_exponent' | 2 | 45 | 1.285241 | 0.286541 | |
| Post-hoc contrasts | B | SE | df | t | p |
| pre_peak - pre_trough | X | X | X | X | X |
| pre_peak - sham | X | X | X | X | X |
| pre_trough - sham | X | X | X | X | X |
| | Df | Df.res | F | p | |
| occipital_theta_abundance' | 2 | 45 | 0.688778 | 0.507405 | |
| Post-hoc contrasts | B | SE | df | t | p |
| pre_peak - pre_trough | X | X | X | X | X |
| pre_peak - sham | X | X | X | X | X |
| pre_trough - sham | X | X | X | X | X |
| | Df | Df.res | F | p | |
| 'occipital_alpha_abundance' | 2 | 45 | 0.014586 | 0.985524 | |
| Post-hoc contrasts | B | SE | df | t | p |
| pre_peak - pre_trough | X | X | X | X | X |
| pre_peak - sham | X | X | X | X | X |
| pre_trough - sham | X | X | X | X | X |

| | Df | Df.res | F | p | |
|---|---|---|---|---|---|
| 'occipital_sigma_abundance' | 2 | 45 | 1.281194 | 0.28764 | |
| Post-hoc contrasts | B | SE | df | t | p |
| pre_peak - pre_trough | X | X | X | X | X |
| pre_peak - sham | X | X | X | X | X |
| pre_trough - sham | X | X | X | X | X |
